# Acceleration and Velocity Dissociate Temporal Phases of Postural Control in Rhesus Macaques

**DOI:** 10.64898/2026.03.20.713289

**Authors:** Olivia M.E. Leavitt Brown, Bassil A. Ramadan, Kathleen E. Cullen

**Affiliations:** Department of Biomedical Engineering, Johns Hopkins University, Baltimore, MD, 21205; Department of Neuroscience, Department of Otolaryngology-Head and Neck Surgery, Kavli Neuroscience Discovery Institute, Johns Hopkins University, Baltimore, MD, 21205

## Abstract

Maintaining balance requires the nervous system to transform sensory signals about unexpected postural perturbations into precisely timed motor commands. Although human studies have established that postural responses unfold in distinct temporal phases, how specific kinematic variables structure these phases during rotational perturbations remains unresolved, because angular acceleration and velocity are typically confounded. Here, we developed a rhesus macaque model of postural control that independently manipulates angular acceleration and peak velocity during transient pitch and roll tilts in monkeys of either sex. By simultaneously measuring head kinematics—directly relevant to vestibular signaling—and center-of-pressure dynamics, we quantified how sensory inputs and motor outputs evolve across successive phases of the postural response. We show that short-latency postural responses (<100 ms) are primarily governed by angular acceleration, whereas medium-latency responses (100–200 ms) scale with angular velocity. This dissociation was robust across perturbation axes and accompanied by axis-dependent control strategies: roll tilts elicited constrained head motion consistent with active stabilization in space, whereas pitch tilts produced more compliant, platform-following behavior. Together, these findings identify distinct kinematic variables governing successive phases of balance control and establish a primate framework for linking neural circuit activity to the temporal organization of postural responses.

**Significance Statement:** Maintaining balance requires transforming sensory signals about unexpected body motion into precisely timed motor commands. Progress in understanding this process has been limited because angular acceleration and velocity are inherently coupled during rotational perturbations. Here, using a rhesus macaque model, we dissociate these kinematic variables and show that they govern distinct temporal phases of postural control: angular acceleration determines short-latency (<100 ms) responses, whereas angular velocity shapes medium-latency (100–200 ms) adjustments. We further demonstrate axis-dependent postural strategies that parallel those observed in humans. Together, these findings resolve a longstanding confound in balance research and establish a primate framework that will enable future studies to link neural circuit activity to the biomechanics of postural control.

## Introduction

Maintaining stable posture in a dynamic environment is essential for everyday activities, and failures of postural stability are a leading cause of falls and mobility impairments (Kakara et al., 2024; Moreland et al., 2023). Stability depends on the integration of vestibular, visual, and proprioceptive inputs to generate coordinated reflexive and voluntary motor responses (Forbes et al., 2020; Mergner et al., 2009). From a neural systems perspective, this requires transforming multisensory information about unexpected perturbations to the body into motor commands that are precisely timed across distinct temporal phases. However, how specific kinematic variables shape these phases during rotational perturbations remains unresolved. We therefore tested whether angular acceleration and velocity selectively govern early and later phases of rotational postural responses.

Human studies of translational perturbations have established a temporal organization of postural responses, with short-latency components dominated by somatosensory inputs and medium-latency components shaped by vestibular signals (Allum et al., 2003; Diener et al., 1988; Forbes et al., 2020; Horak et al., 1990; Inglis & Macpherson, 1995). In these paradigms, acceleration and velocity were historically confounded, but later work demonstrated that acceleration primarily governs short-latency responses, whereas velocity shapes medium-latency adjustments (Welch & Ting, 2009). Whether a similar dissociation governs postural control during rotational perturbations—where semicircular canal signals are central—remains unknown.

Rotational perturbations provide a sensitive assay for probing vestibular contributions to balance, as responses are disproportionately impaired following vestibular loss (cat: Inglis & Macpherson, 1995; human: Horak et al., 2016). Prior studies have further revealed axis-dependent strategies: roll tilts typically elicit symmetric stabilizing responses, whereas pitch tilts often produce forward–backward asymmetries and greater platform-following motion (Allum et al., 2008; Carpenter et al., 2001; Horak et al., 2016; Mansfield & Maki, 2009; Welch & Ting, 2009). However, in all prior rotational studies, angular acceleration and velocity have remained inherently coupled, preventing independent assessment of their contributions to successive temporal phases of the postural response.

Accordingly, here we developed a rhesus macaque model of postural control that independently manipulates angular acceleration and peak velocity during transient pitch and roll tilts. Rhesus macaques share key biomechanical features with humans, including plantigrade stance and comparable base-of-support geometry, while permitting direct recordings from and manipulations of neural circuits that are not feasible in human subjects. At the same time, interspecies differences, including variations in center-of-mass height relative to limb length, habitual postural configuration, and base-of-support geometry, preclude direct extrapolation from human data and necessitate species-specific behavioral benchmarks to enable meaningful interpretation of future neurophysiological findings. Providing benchmarks for rhesus monkey postural control within this framework enables future neurophysiological investigations—combining single-neuron recordings and causal circuit manipulations—that can directly link neural circuit activity to postural control. By disentangling acceleration and velocity, we directly test how the nervous system parses unexpected postural perturbations into temporally distinct control signals. Because head motion both drives vestibular afferent activity and reflects postural motor output within a closed-loop system, we simultaneously quantified six-dimensional head kinematics together with center-of-pressure dynamics. Our findings show that short-latency postural responses (0-100 ms) are primarily governed by angular acceleration, whereas medium-latency responses (100–200 ms) scale with angular velocity. This dissociation extends principles previously established for translational perturbations to rotational postural control. In addition, roll tilts elicited constrained head motion consistent with enhanced stabilizing control in space, whereas pitch tilts produced more compliant, platform-following behavior, revealing axis-dependent postural strategies. Together, our findings identify distinct kinematic variables that structure successive phases of balance control and establish a primate framework for linking neural circuit activity to the temporal organization and control variables of postural responses.

## Methods

Three rhesus monkeys (Macaca mulatta; two male, ‘Monkey E’ and ‘Monkey B,’ and one female, ‘Monkey D’) were used in this study. Each animal completed ≥20 trials per perturbation condition across multiple stimulus conditions. All experimental procedures were approved by the Johns Hopkins Animal Care and Use Committee (ACUC, protocol number PR22M342). The ACUC is accredited by the American Association for the Accreditation of Laboratory Animal Care (AAALAC) and is in compliance with the guidelines of the Office of Laboratory Animal Welfare at the National Institutes of Health. As described previously (Zobeiri & Cullen, 2022), each animal was anesthetized and equipped with a titanium post fastened to the skull using titanium screws and dental acrylic to allow immobilization of the head and securing of recording hardware. All animals recovered for at least two weeks before experiments began.

### Experimental Setup

The experimental setup consisted of a hexapod motion platform (SYMETRIE, Nimes, France) onto which was mounted a custom-built, clear-sided behavioral chamber with a fiberglass frame. The behavioral chamber is a total of 2.5m in length to accommodate walking experiments, but Plexiglas dividers allow for confinement of the animal to any of three sections. The chamber was 0.5m wide and 0.75 m high, allowing animals of any size to sit, stand, walk several strides, and turn around. All experiments reported here took place with the animal confined to the central section of the chamber, which was 0.7 m long and was also instrumented with a force plate. The custom-built force plate consisted of four ABS plastic footplates each mounted to a uniaxial load cell (Omega Engineering, Sunbury, OH), all of which were mounted to a single baseplate. Animals were transported from the colony room to the setup in a standard primate chair, then fitted with a wireless, wearable inertial measurement unit (IMU; Shimmer Sensing, Dublin, Ireland) and 3D-printed optical orientation tracker (Vagvolgyi et al., 2022) affixed to their head fixation post. Additionally, four high-speed cameras (Teledyne FLIR, Wilsonville, OR) were mounted around the setup to capture video of the animals, which was used post hoc to confirm that postural responses were free of step responses.

### Perturbation Design

Perturbations were explicitly designed to dissociate angular acceleration from peak velocity while maintaining comparable total displacement. Perturbations were designed in MATLAB by first constructing an angular acceleration time series which included an acceleration and deceleration block with the desired peak acceleration amplitude, and duration and inter-block interval that together yielded the desired peak velocity and total displacement. The acceleration blocks were constructed to mimic, as closely as possible, a step change in acceleration given the temporal resolution of the hexapod motion platform (100 samples/sec). This acceleration signal was then double-integrated, and the resulting position signal was fed into the motion platform.

Perturbations were based on those used in similar previous experiments in humans (Allum et al., 2008; Carpenter et al., 2001) and cats (Inglis & Macpherson, 1995), which had a peak velocity of 40 deg/s and an estimated angular acceleration of 500 deg/s^2^. These values were used as the standard “base” perturbation, which was shared between the set varying angular acceleration, and the set varying peak velocity. For each set (velocity and acceleration) a higher and lower value than the base perturbation was used: 200 and 1000 deg/s^2^ accelerations, and 20 and 60 deg/s peak velocities. This yielded a total of five perturbations, which were delivered in four directions: forward and backward pitch, and leftward and rightward roll. Overall, there were 20 perturbation conditions.

### Conceptual Biomechanical Model

To provide mechanistic intuition for how passive mechanical properties can shape head and body motion during platform tilts, we implemented a conceptual, simplified dual-pendulum biomechanical model. The model represents the body as two rigid segments: a trunk–limb segment pivoting about the ankles and a head segment articulated at the neck.

Platform perturbations were imposed by prescribing the angular velocity of the support surface as a time-varying input, mirroring the experimental perturbation design. This velocity signal was numerically integrated to obtain platform angular displacement. The equations of motion included linear stiffness and damping terms coupling the trunk to the rotating platform, as well as stiffness and damping terms coupling the head to the trunk (see Supplementary Table S1).

Segment masses, lengths, and moments of inertia were set to representative values based on standard rhesus monkey biometric tables (Vilensky, 1979). Because neck and body stiffness have not been directly measured in monkeys, stiffness and damping parameters were chosen within physiologically plausible ranges reported for cats and humans (Goldberg & Peterson, 1986; Keshner et al., 1992; Peng et al., 1996; Ting & Macpherson, 2004). Importantly, these parameters were not tuned to fit the experimental data but were selected solely to illustrate qualitatively distinct regimes of platform-following versus head-in-space stabilization, rather than to reproduce the experimental responses.

Torques on the trunk and head were computed using the following equations, where θ₁ and θ₂ denote trunk and head angles relative to vertical, respectively, φ denotes platform angle, and overdots indicate time derivatives:

Torque on trunk (τ₁):

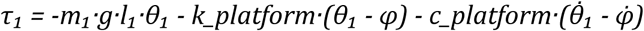

Torque on head (τ₂):

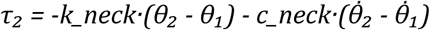

Here, *m₁*, *l₁*, and *I₁* denote trunk mass, center-of-mass distance, and moment of inertia, respectively; *I₂* denotes head moment of inertia; and k and c terms represent stiffness and damping coefficients.

Angular accelerations were computed as:

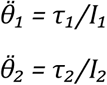

Head and trunk angles and angular velocities were obtained by numerically integrating the resulting system of ordinary differential equations using MATLAB’s ODE45 solver.

### Behavioral Training & Measurement

As stated in the description of the behavioral setup, the behavioral chamber was instrumented with a force plate, and the animal wore a wireless IMU and an optical tracker secured to its headpost during training and experimental sessions. Each monkey was acclimated to the behavioral chamber over the course of three training sessions, which took approximately one week. During the first session, the monkey was repeatedly transferred between the transport chair and the behavioral chamber and rewarded with food treats following each transfer attempt. During the second session, the animal was allowed to enter the central chamber and freely explore the space with intermittent juice rewards from a spout fixed to the ceiling of the chamber; animals quickly learned to orient themselves toward the reward spout. During the third session, the animal was again transferred to the chamber, and sinusoidal platform motions were applied to acclimate the animal to platform movement, minimizing startle responses during experiments. Each of these behavioral training sessions took between 30 minutes and 2 hours. The longest training session exceeded the duration of most experimental sessions, ensuring acclimation to extended periods in the setup.

Experimental sessions began after the animal had completed training sessions and reinforced the trained standing behavior throughout testing. For each experimental session, the animal was first transferred to the chamber and stood on the force plate. Data from the force plate were continuously fed to a real-time experimental control computer, and the force distribution across each of the four panels was calculated. Head orientation data from the optical head tracker were fed into the same system. To ensure consistent initial stance, the animal was required to stand symmetrically at trial onset. The criteria for starting a trial included having one paw on each footplate with weight balanced across the four footplates (i.e., <10 N difference between the front and back halves and between the left and right halves of the force plate). The animal also needed to have its head vertical and facing straight ahead (deviations <10° in any axis). Transient trials began at random times, occurred in a randomized order, and shared identical total displacement to minimize anticipatory postural responses. Before each trial, the experimenter initiated recording on the motion capture cameras, while the force plate and IMU recorded continuously at 1,000 samples/s. Data collection sessions lasted as long as the animal remained cooperative with the experiment. For each monkey, a minimum of 20 trials were collected per condition.

### Data Analyses

Following experiments, data were calibrated and synchronized in MATLAB using a custom code pipeline. Data from the load cells under each footplate of the force plate were converted to center of pressure (CoP) position according to the following equations:

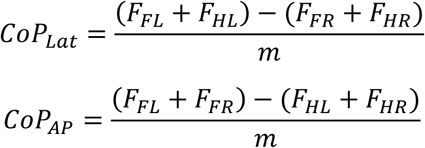

where *F_FL_*, *F_FR_*, *F_HL_*, and *F_HR_* represent the forces on the front left, front right, hind left, and hind right plates, respectively, and *m* represents the mass of the monkey. Data from the IMUs were calibrated onboard and synchronized to the same clock onboard each device.

Camera data were fed through DeepLabCut and Anipose (Karashchuk et al., 2021; Mathis et al., 2018) for markerless pose reconstruction. This video-based data was primarily used to determine whether step responses occurred during a trial; trials containing step responses were excluded from further analysis. Trials were also removed by the program if head velocity in the 500 ms preceding a trial exceeded 20 deg/s, and if head velocity at any point in the trial (including 500 ms before the onset and after the offset of motion) exceeded 400 deg/s, which indicated head shaking or other voluntary movement contaminating the animal’s postural response. Approximately 10% of trials collected for each monkey were excluded based on video, kinematic, or kinetic data. This rate of exclusion was comparable between animals.

Trials of the same perturbation in the same animal were then synchronized, and the mean response and standard error were computed. We then computed the mean response across the first 100 ms following perturbation onset (short-latency window) and during the 100–200 ms interval (medium-latency window). Response onset latency was defined as the first time point at which the response exceeded the root mean square (RMS) error computed over the 500 ms preceding perturbation onset. Total angular displacement was computed as the integral of angular velocity, total linear displacement as the double integral of acceleration, and CoP trajectory length as the cumulative sum of the CoP response (Lemay et al., 2014). All displacement measures were computed over the entire duration of the platform motion.

Finally, to quantify symmetry between responses to perturbations in opposite directions (left vs. right, forward vs. backward), we computed an “asymmetry index” (adapted from Kong et al., 2010). For the within-direction comparison, we calculated the distribution of the mean absolute percentage error (MAPE) across all possible pairs of responses to the same stimulus within a given direction (e.g., all leftward responses, all rightward responses). The resulting distributions for the two opposing directions were then combined to yield the first distribution. For the across-direction comparison, we inverted the responses for one direction and then calculated the MAPE across all possible pairs of responses regardless of direction, producing a second distribution. The asymmetry index was defined as the difference between the mean values of these two distributions.

### Statistical Analyses

Statistically significant differences between monkeys and between conditions within the velocity and acceleration sets were established using a two-way ANOVA accounting for unequal sample sizes. A significance threshold of p < 0.025 was used for differences between monkeys and differences between conditions to control for multiple comparisons across factors. When a significant main effect was detected in the ANOVA, linear regression was used to quantify the dependence of the pooled responses across all monkeys on angular acceleration or peak velocity, as appropriate. Significance of the asymmetry index was assessed using a two-tailed t-test (α = 0.05), with the null hypothesis of symmetry (no difference between distributions) and the alternative hypothesis of asymmetry. For each perturbation axis, p-values across monkeys and stimuli were then combined using Fisher’s method.

## Results

In this study, we examined the postural responses of rhesus monkeys to tilt perturbations on a hexapod motion platform (**Fig. 1A**). Our goal was to establish a nonhuman primate model of postural control and determine how different kinematic variables—axis, angular velocity, and acceleration—influence postural strategies. Three monkeys were trained to stand symmetrically in a natural perching posture on a force plate, with their heads level and facing forward. Each animal was outfitted with a head-mounted inertial measurement unit (IMU) and an optical tracking marker (Vagvolgyi et al., 2022) to ensure initial head alignment was consistent across trials. Once trained, we applied tilt perturbations and recorded the resulting head motion and center-of-pressure (CoP) responses. Tilt perturbations were applied in both pitch (forward/backward) and roll (left/right) axes (**Fig. 1B**), with peak velocity and angular acceleration varied independently to isolate their specific contributions (**Fig. 1C**). High-speed video from four cameras was analyzed using 3D markerless pose estimation (i.e., DeepLabCut and Anipose, (Karashchuk et al., 2021; Mathis et al., 2018)) to confirm the absence of stepping or non-postural voluntary movements such as head shaking. **Figure 1D** illustrates a simple dual-pendulum model used here as a conceptual framework to distinguish passive mechanical responses from active stabilization. Varying body–platform coupling and neck stiffness produces qualitatively distinct head and body kinematics, providing intuition for how different postural strategies—ranging from platform-following to stabilization in space—can arise even in the absence of active neural control.

**Figure 1.**
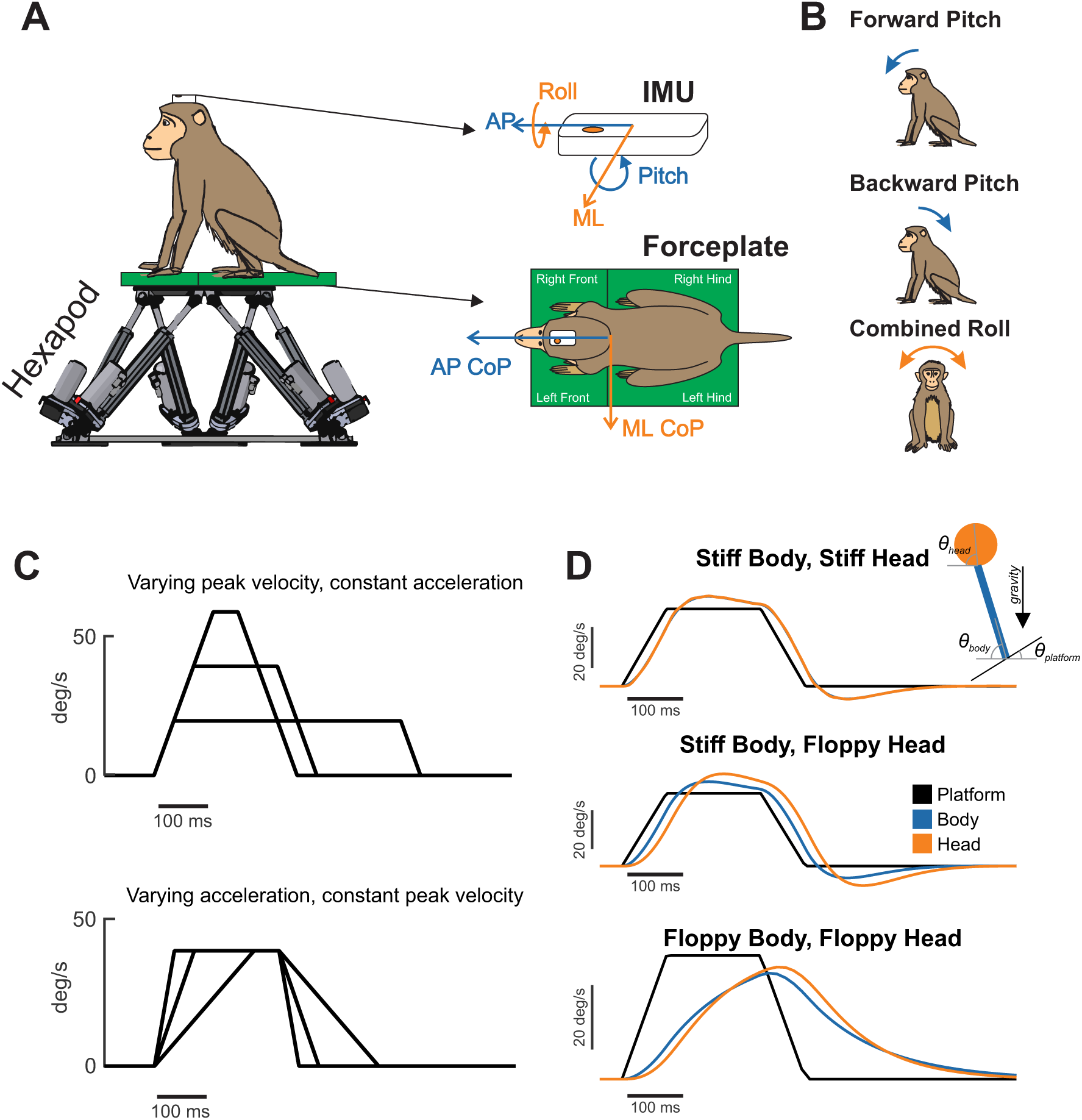
Experimental paradigm and conceptual framework. (A) Rhesus monkeys stood freely on a force plate mounted to a hexapod motion platform while head kinematics were measured using an inertial measurement unit (IMU). Center-of-pressure (CoP) was computed from ground-reaction forces recorded beneath each limb. (B) Transient rotational perturbations were applied in pitch (forward/backward) and roll (left/right). (C) Perturbation profiles were designed to independently vary peak velocity while holding angular acceleration constant (top), or vary angular acceleration while holding peak velocity constant (bottom). (D) A simple dual-pendulum model (see Methods) illustrating how variation in body–platform coupling and neck stiffness can produce distinct head and body kinematics, providing a conceptual framework for distinguishing platform-following behavior from active stabilization.

## Modulation of Pitch Postural Responses by Stimulus Velocity

Most prior studies of postural control have focused on rotational perturbations in the pitch axis (reviewed in Forbes et al., 2020). Thus, we began by examining responses to pitch tilts to specifically address how postural responses vary as a function of peak velocity. Importantly, by holding angular acceleration constant, we were able to isolate velocity-specific effects. We first applied transient forward pitch tilts (of 20, 40, or 60 deg/s) with a fixed angular acceleration (500 deg/s²) - parameter values based on prior work in quadrupedal cats (Macpherson et al., 2007; **Fig. 1B**). Mean head velocity, acceleration, and CoP (center of pressure) along the perturbation axis are shown for each monkey in **Figure 2A** (Monkey D) and **Figure S1A, B** (Monkey E & Monkey B).

**Figure 2.**
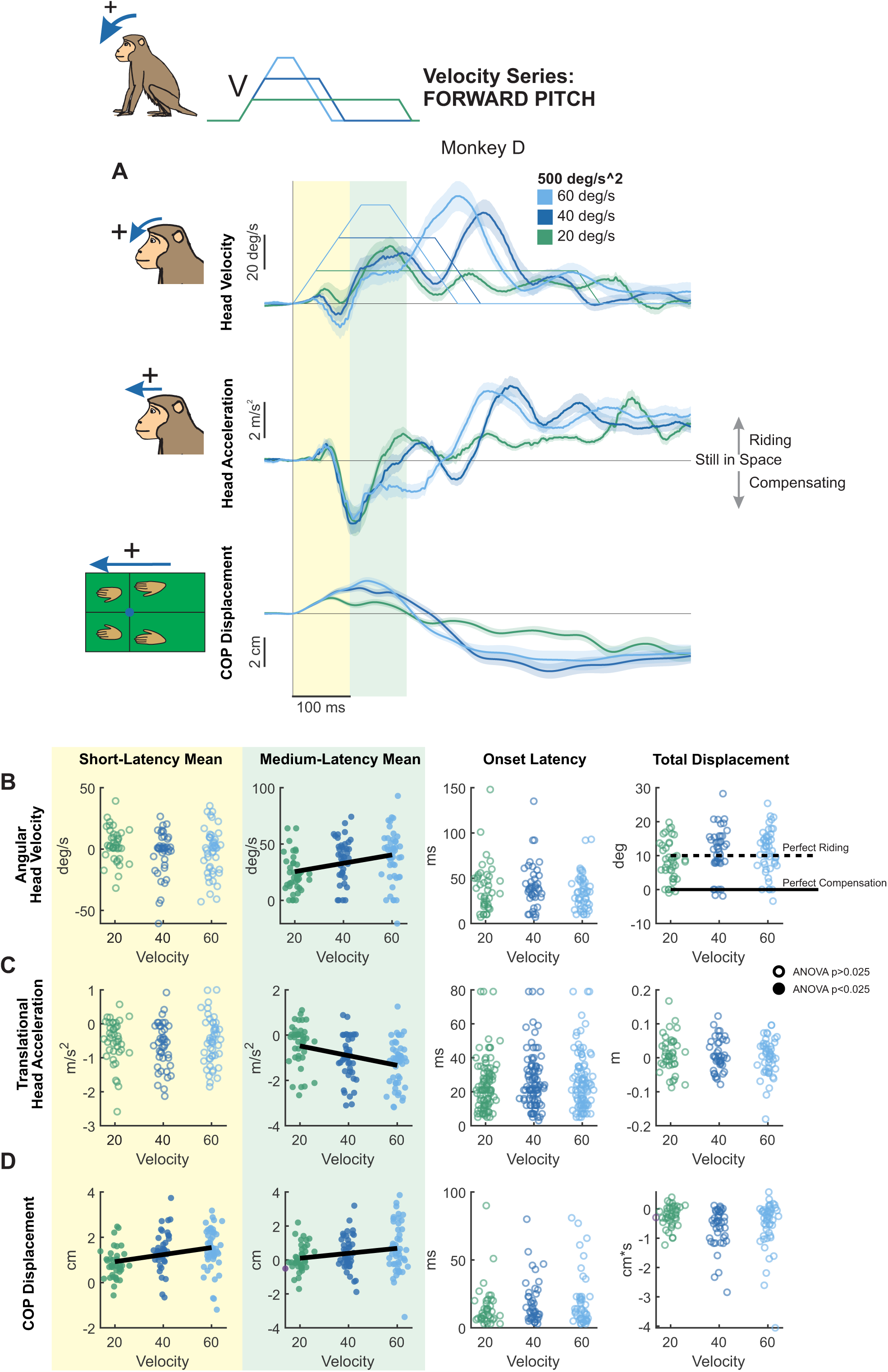
Postural responses to forward pitch perturbations of varying peak velocity in the axis of the perturbation. (A) Head pitch velocity, head fore-aft acceleration, and fore-aft CoP responses of “Monkey D” to perturbations with peak velocities of 20 (purple), 40 (blue), and 60 (green) deg/s and acceleration of 500 deg/s^2^. Traces show the mean response over 20 trials and shaded regions represent the mean +/- SEM. (B) Quantified head pitch velocity response. Dots and open circles represent individual trials, pooled across monkeys. Filled circles indicate statistical significance (ANOVA p<0.025) for comparison across velocities accounting for differences between individual monkeys. Open circles indicate lack of statistical significance (ANOVA p>0.025). Bold black lines indicate a linear velocity-dependent trend (regression p<0.05). From left to right: Yellow region: mean head pitch velocity in the short-latency window (ANOVA p=0.59). Green region: mean head pitch velocity in the medium-latency window (ANOVA p=0.000943, regression p=0.000543). White background, left: response onset latency (ANOVA p=0.35). White background, right: total head pitch displacement (ANOVA p=0.050372). Dotted black line indicates perfect riding response, solid black line indicates perfect compensatory response leading to no head-in-space motion. See Figure S1 for mean responses of individual monkeys. (C) Quantified head fore-aft acceleration response, same format as (B). From left to right, mean acceleration in the short-latency window (ANOVA p=0.61), mean acceleration in the medium-latency window (ANOVA p=0.000615, regression p=6.26e-5), response onset latency (ANOVA p=0.53), and linear displacement (ANOVA p=0.68227). (D) Quantified CoP fore-aft response, same format as (B) and (C) From left to right, mean CoP response in the short-latency window (ANOVA p=0.0016, regression p=0.000572), mean CoP response in the medium-latency window (ANOVA p=0.0202, regression p=0.0163), CoP response onset latency (ANOVA p=0.96), and CoP distance traversed (ANOVA p=0.0350).

Overall, we found that pitch perturbations reliably elicited head movements in monkeys that closely resembled those previously reported in cats (Macpherson et al., 2007; Torres-Oviedo et al., 2006). These head motion responses were relatively consistent across animals and showed low trial-to-trial variability, particularly 100 ms after the onset of the perturbation. To quantify their temporal dynamics, we defined two windows of interest based on the prior literature: a short-latency window (0–100 ms) and a medium-latency window (100–200 ms) (**Fig. 2A** yellow and green shaded regions, respectively; Allum et al., 2003; Diener et al., 1988; Forbes et al., 2020; Horak et al., 1990; Inglis & Macpherson, 1995). Within each window, we then computed the mean head pitch velocity, fore-aft acceleration, and CoP response. **Figures. 2B-D** plot the results of this analysis, for short-latency (yellow), medium-latency (green), onset latency (white, left), and distance traversed (white, right) measurements for each of the three transient perturbation profiles (i.e., peak velocities reaching 20, 40, or 60 deg/s). The results for each individual monkey are provided in **Fig. S1C-E**.

### Head motion responses

Our analysis of head pitch velocity revealed that, in the short-latency window (**Fig. 2B**, yellow panel), mean head pitch velocity remained relatively constant across velocities. In contrast, during the medium-latency window (**Fig. 2B**, green background), pitch velocity increased systematically with stimulus velocity, demonstrating significant velocity-dependent modulation. Correspondingly, our analysis of fore-aft translational head acceleration revealed a similar pattern (**Fig. 2C**); medium-latency responses decreased significantly, increasing in magnitude as a function of stimulus velocity (**Fig. 2C**, green background), whereas this was not the case for short-latency acceleration responses (**Fig. 2C**, yellow panel). Onset latencies of head pitch velocity and fore-aft translational acceleration were comparable (largely within 50ms) and did not differ significantly across velocity conditions (**Fig. 2B,C** left white panels), indicating that response onset was independent of peak stimulus velocity. Furthermore, total head pitch displacement was 10 degrees in each case, indicating that the monkey rode on the platform over the course of the perturbation (**Fig 2B**, right white panel). Unless otherwise noted, onset latencies and total displacements of head and CoP motion were invariant across conditions. Together, these results indicate that rhesus monkey head motion responses to pitch tilts exhibit significant velocity-dependent modulation, which emerges primarily during the medium-latency phase.

### CoP measures

We next assessed fore-aft CoP displacement as a dynamic indicator of whole-body stabilization to pitch perturbations (**Fig. 2D**). In contrast to head motion kinematics, CoP responses demonstrated significant velocity dependence during the short-latency window (**Fig. 2D**, left panel) as well as the medium-latency window (**Fig. 2D**, green panel), with both increasing significantly as a function of velocity. Qualitative inspection of the CoP traces in Fig 2A further revealed that the peak CoP response was coincident with the peak velocity of the platform perturbation. This time point occurred within the short-latency window during the 20 deg/s condition and the medium-latency window during the 40 and 60 deg/s conditions, suggesting that velocity-scaled adjustments may occur coincident with this transition point. Taken together, these results demonstrate that pitch perturbations generally elicited a passive “ride-the-platform” response (Buchanan & Horak, 2001) – where the animal’s head and body moved along with the platform rather than stabilized relative to space, with velocity modulating medium-latency adjustments. Within the conceptual framework illustrated in **Fig. 1D**, this pattern qualitatively resembles a regime in which the body is strongly coupled to the platform while the neck remains compliant, resulting in platform-following motion of the trunk and lagged head motion.

### Direction-Dependent Modulation

We next quantified responses to backward tilts. **Figure 3** shows the same analyses applied to forward tilts (**Figure 2**) but for backward tilts with the same fixed angular acceleration (500 deg/s²) and peak velocities of 20, 40, or 60 deg/s. In contrast to forward tilts, responses to backward perturbations were less stereotyped across animals (compare **Fig. 3A** and **Fig. S2A,B**). Correspondingly, there were significant differences in mean velocity between monkeys during both the short and medium-latency window (**Fig. S2C**). On average, pitch velocity in the short-latency window remained unchanged across stimulus velocities (**Fig. 3B**, yellow panel), whereas it significantly increased in magnitude as a function of stimulus velocity (i.e., continued to move along with platform as speed increased) in the medium-latency window (**Fig. 3B**, green panel). Interestingly, at the lowest peak velocity head pitch displacement was around 10 degrees, demonstrating a riding-the-platform response. Total displacement also decreased significantly with increasing velocity, indicating greater compensation as velocity increased (**Fig. 3B**, right white panel). Head acceleration responses also followed this pattern: Mean acceleration in the short-latency window did not vary with stimulus velocity (**Fig. 3C**, yellow panel), but increased significantly in the medium-latency window (**Fig. 3C**, green panel).

**Figure 3.**
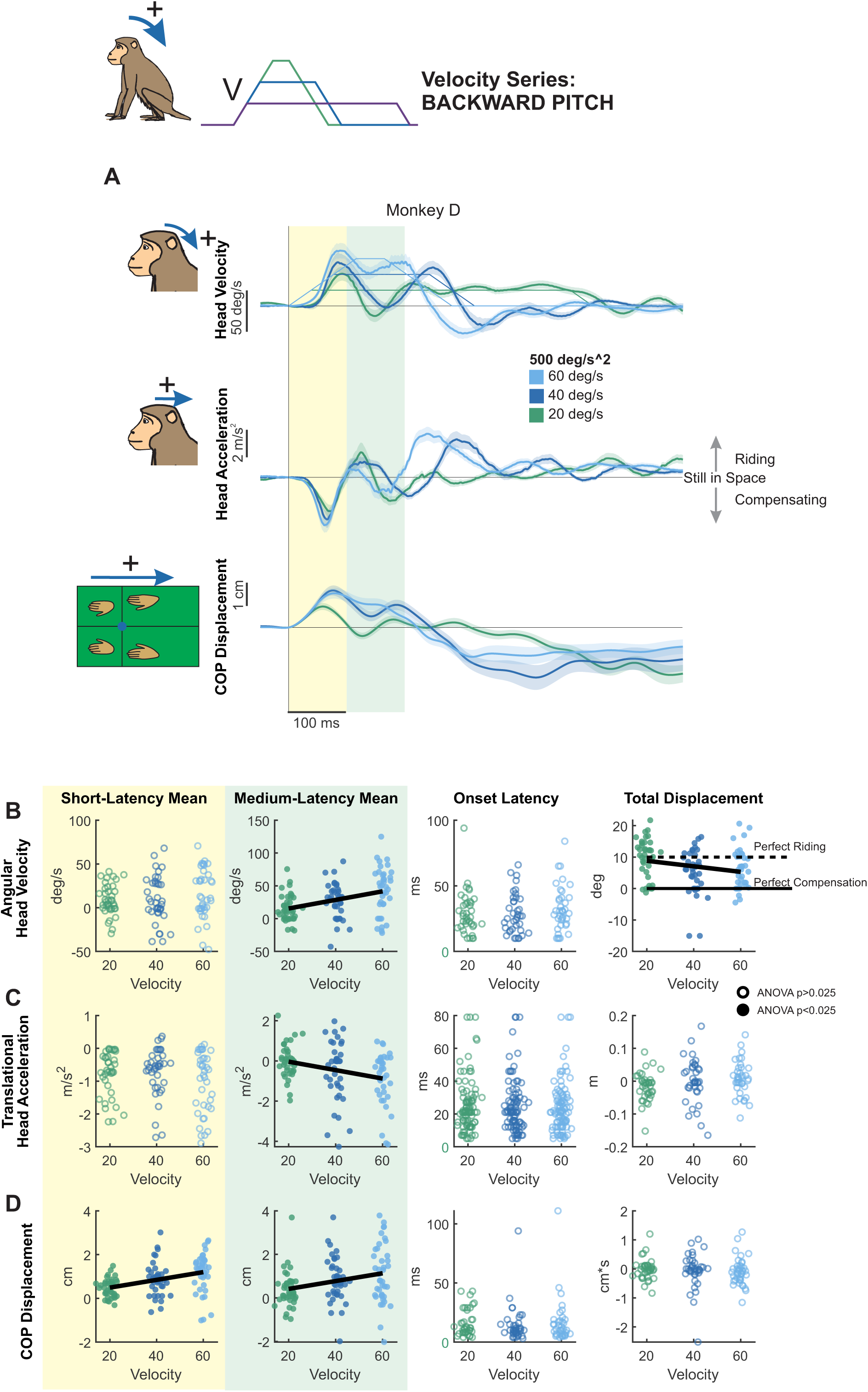
Postural responses to backward pitch perturbations of varying peak velocity in the axis of the perturbation. (A) Head pitch velocity, head fore-aft acceleration, and fore-aft CoP responses of “Monkey D” to perturbations with peak velocities of 20 (purple), 40 (blue), and 60 (green) deg/s and acceleration of 500 deg/s^2^. Traces show the mean response over 20 trials and shaded regions represent the mean +/- SEM. (B) Quantified head pitch velocity response, same plotting conventions as Figure 2. From left to right: Yellow region: mean head pitch velocity in the short-latency window (ANOVA p=0.64). Green region: mean head pitch velocity in the medium-latency window (ANOVA p=0.000218, regression p=8.70e-5). White background, left: response onset latency (ANOVA p=0.45). White background, right: total head pitch displacement (ANOVA p=0.00319, regression p=0.0203). Dotted black line indicates perfect riding response, solid black line indicates perfect compensatory response leading to no head-in-space motion. See Figure S2 for mean responses of individual monkeys. (C) Quantified head fore-aft acceleration response, same format as (B). From left to right, mean acceleration in the short-latency window (ANOVA p=0.41), mean acceleration in the medium-latency window (ANOVA p=2.24e-5, regression p=0.00344), response onset latency (ANOVA p=0.92) linear displacement (ANOVA p=0.23). (D) Quantified CoP fore-aft response, same format as (B) and (C) From left to right, mean CoP response in the short-latency window (ANOVA p=0.000189, regression p=1.77e-5), mean CoP response in the medium-latency window (ANOVA p=0.00386, regression p=0.00369), CoP response onset latency (ANOVA p=0.84) CoP distance traversed (ANOVA p=0.86).

Lastly, while head movement responses were less stereotyped across monkeys for backward versus forward pitch perturbations, quantification of CoP revealed similarities across monkeys (**Fig. 3D, Fig. S2E**). As for forward tilts, CoP responses to backward tilts exhibited significant velocity-related effects during both the short- and middle-latency windows (**Fig. 3D**, yellow and green panels), increasing in magnitude as a function of increasing velocity. Thus, like forward tilts, CoP responses to backward tilts were modulated by stimulus velocity even during the short-latency phase. Further, similar to forward tilts, these results indicate that backward tilt perturbations generally elicit a passive “ride-the-platform” response at low velocity, with increasing compensation at higher velocities.

## Modulation of Roll Postural Responses by Stimulus Velocity

We next applied tilt perturbations in the roll axis, again varying peak velocity independently of acceleration (**Figure 4**). In contrast to the markedly asymmetrical response to forward versus backward pitch tilts described above, responses to leftward and rightward roll tilts were relatively symmetrical across animals (compare **Fig. S3 A** versus **B**). We quantified this observation by computing an asymmetry index for leftward versus rightward, and forward versus backward tilt perturbations (see Methods). Overall, we found that roll responses were consistently symmetrical across monkeys, while pitch responses were a mix of symmetrical and asymmetrical. Given that roll perturbation response were relatively symmetric, we inverted responses to rightward perturbations and combined them with those from leftward perturbations in our analysis, resulting in a total of 40 trials per condition.

**Figure 4.**
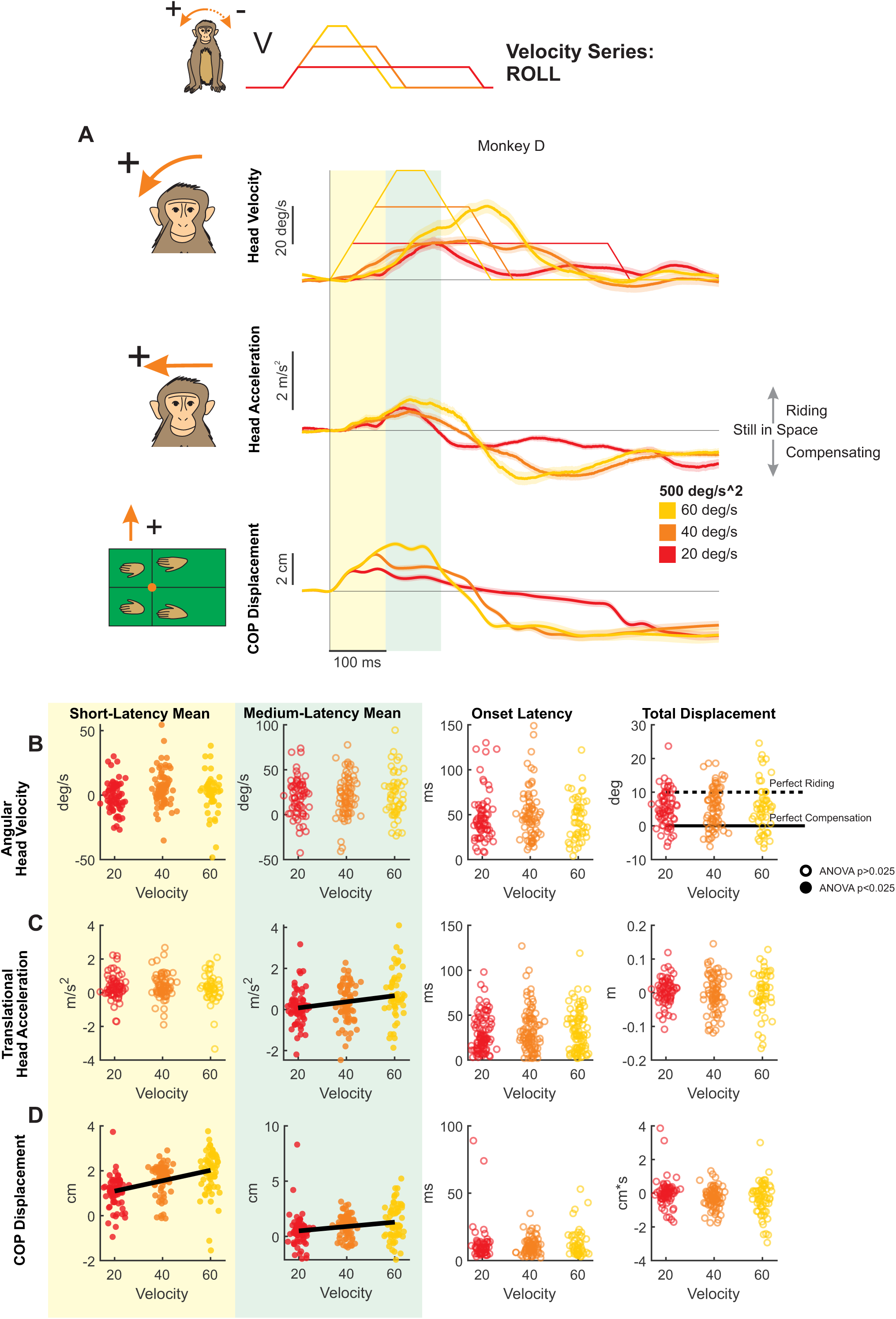
Postural responses to roll perturbations of varying peak velocity in the axis of the perturbation. (A) Head roll velocity, head lateral acceleration, and lateral CoP responses of “Monkey D” to perturbations with peak velocities of 20 (red), 40 (orange), and 60 (yellow) deg/s and acceleration of 500 deg/s^2^. Traces show the mean response over 20 trials and shaded regions represent the mean +/- SEM. (B) Quantified head roll velocity response, same plotting conventions as Figure 2. From left to right: Yellow region: mean head roll velocity in the short-latency window (ANOVA p=0.0104). Green region: mean head roll velocity in the medium-latency window (ANOVA p=0.0210, regression p=0.00883). White background, left: response onset latency (ANOVA p=0.59). White background, right: total head roll displacement (ANOVA p=0.79). Dotted black line indicates perfect riding response, solid black line indicates perfect compensatory response leading to no head-in-space motion. See Figure S4 for mean responses of individual monkeys. (C) Quantified head lateral acceleration response, same format as (B). From left to right, mean acceleration in the short-latency window (ANOVA p=0.60), mean acceleration in the medium-latency window (ANOVA p=0.00195, regression p=0.00107), response onset latency (ANOVA p=0.47) linear displacement (ANOVA p=0.76). (D) Quantified CoP lateral response, same format as (B) and (C) From left to right, mean CoP response in the short-latency window (ANOVA p=1.81e-11, regression p=2.54e-11), mean CoP response in the medium-latency window (ANOVA p=0.000533, regression p=0.000608), CoP response onset latency (ANOVA p=0.39) CoP distance traversed (ANOVA p=0.0448).

Notably, unlike pitch perturbations, roll tilts revealed a qualitatively distinct postural strategy, characterized by constrained head motion suggestive of active stabilization. Overall, individual monkeys generated more consistent responses to roll than pitch perturbations (e.g., compare **Fig. 4A** versus **Fig. 2A** & **Fig. 3A**). Moreover, whereas monkeys tended to ride the platform during pitch tilts, head motion during roll perturbations appeared to be actively constrained—not exceeding the maximum platform velocity of 60 deg/s, and in the case of Monkey D, never exceeding the peak platform velocity of the condition. This result suggests a fundamentally different postural strategy in the roll axis, emphasizing active stabilization over passive compliance. Within the conceptual framework illustrated in Fig. 1D, this behavior falls outside the range expected from passive body–platform mechanics and instead corresponds to a regime requiring active stabilization to constrain head motion relative to space.

To next quantify how stimulus velocity shaped this active stabilization strategy, we measured head roll and lateral head movement and CoP responses in short and medium-latency windows (**Fig 4B-D**, individual monkey responses in **Fig. S4**). We then compared these measures across the three perturbation conditions (i.e., 20, 40, or 60 deg/s).

### Head motion

During the short-latency window, mean head roll velocity did not vary linearly as a function of velocity (**Fig. 4B**, yellow). Medium-latency responses, however, showed inter-subject variability, with two monkeys demonstrating oppositely directed trends, and the third monkey showing no velocity-dependent scaling (**Fig. S4**, green). This heterogeneity resulted in no overall effect of acceleration on medium-latency responses (**Fig. 4B**, green). Similarly, mean lateral head acceleration during the short-latency window was unaffected by peak velocity (**Fig. 4C**, yellow). However, as observed for pitch perturbations, mean head acceleration increased significantly with stimulus velocity during the medium-latency window (**Fig. 4C**, middle). Finally, total roll displacement was consistent across velocities and fell within the band between perfect compensation and perfect riding (i.e., between 0 and 10 degrees) for all monkeys, indicating partial compensation for the platform motion (**Fig 4B**, right white).

### CoP measures

Both short-latency (**Fig. 4D**, yellow) and medium-latency CoP responses (**Fig. 4D**, green) increased significantly with peak velocity, thereby paralleling the trends observed above for backward and forward pitch.

Thus, in summary, rhesus monkeys showed axis-dependent postural strategies to transient tilt perturbations. For pitch tilts in both the forward and backward directions, monkeys tended to “ride” the platform, showing minimal compensation, and head motion responses were velocity-dependent in the medium-latency window, while CoP responses showed velocity-dependent effects in both short- and medium-latency phases. In contrast, the smaller head-in-space motion responses to roll tilts indicated active compensation for platform motion, and were less velocity dependent. However, CoP measures again scaled with velocity in both latency windows, which suggests that the active compensation at the support surface which ultimately constrains head motion is modulated by velocity.

## Modulation of Pitch and Roll Postural Responses by Stimulus Acceleration

So far, we have shown that peak velocity predominantly modulates responses in the medium-latency window. We next hypothesized that acceleration—when varied independently—would also shape short-latency responses based on prior studies of translational perturbations (Welch & Ting, 2009). To test this proposal, we applied tilt perturbations at three acceleration levels (200, 500, and 1000 deg/s²) while holding peak velocity constant at 40 deg/s (**Fig. 1C**). Notably, this design included a shared condition (500 deg/s² at 40 deg/s) with the experiments described above focused on isolating the effect of velocity (**Fig. 1B**), enabling a direct comparison.

### Forward pitch

We first examined head responses to acceleration-controlled forward pitch perturbations (**Fig. 5A**). While in two monkeys there was a trend of increasing mean head pitch velocity in the short-latency window (**Fig. S5**, monkeys E and B), overall, quantification of responses revealed that mean head pitch velocity showed significant differences between accelerations but without a linear dependence **(Fig. 5B**, left panel). Likewise, quantification of responses in the medium-latency window mean head velocity increased as a function of acceleration across animals (**Fig. 5B**, middle). In contrast, head fore-aft linear acceleration was not significantly influenced by acceleration in either the short-latency (**Fig. 5C**, left), or medium-latency (**Fig. 5C**, middle) windows.

**Figure 5.**
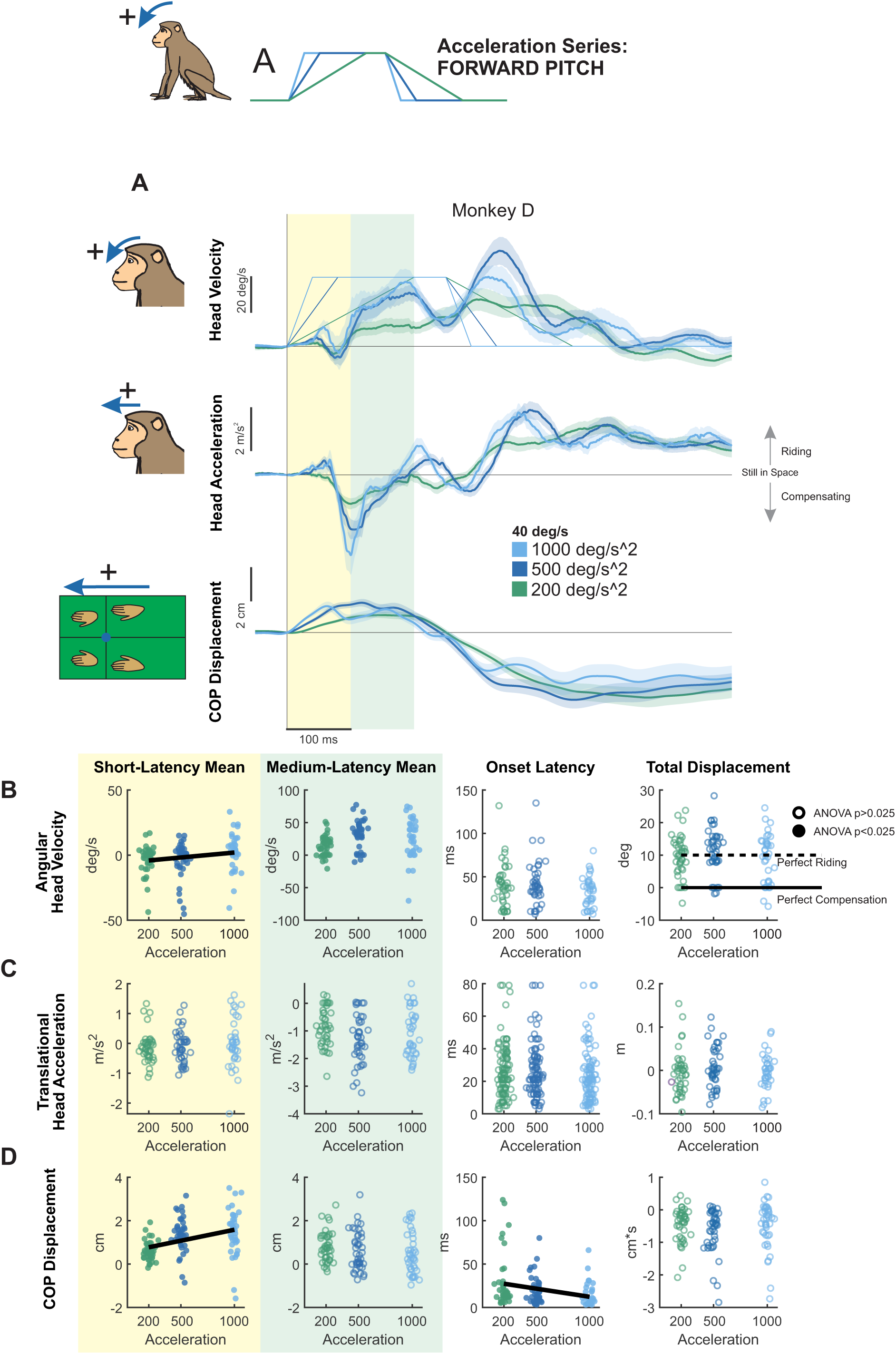
Postural responses to forward pitch perturbations of varying acceleration in the axis of the perturbation. (A) Head pitch velocity, head fore-aft acceleration, and fore-aft CoP responses of “Monkey D” to perturbations with accelerations of 200 (purple), 500 (blue), and 1000 (green) deg/s^2^ and velocity of 40 deg/s. Traces show the mean response over 20 trials and shaded regions represent the mean +/- SEM. (B) Quantified head pitch velocity response. Dots and open circles represent individual trials, pooled across monkeys. Filled circles indicate statistical significance (ANOVA p<0.025) for comparison across velocities accounting for differences between individual monkeys. Open circles indicate lack of statistical significance (ANOVA p>0.025). Bold black lines indicate a linear acceleration-dependent trend (regression p<0.05). From left to right: Yellow region: mean head pitch velocity in the short-latency window (ANOVA p=0.0218, regression p=0.0384). Green region: mean head pitch velocity in the medium-latency window (ANOVA p=0.00204, regression p=0.33). White background, left: response onset latency (ANOVA p=0.17). White background, right: total head pitch displacement (ANOVA p=0.35). Dotted black line indicates perfect riding response, solid black line indicates perfect compensatory response leading to no head-in-space motion. See Figure S5 for mean responses of individual monkeys. (C) Quantified head fore-aft acceleration response, same format as (B). From left to right, mean acceleration in the short-latency window (ANOVA p=0.47), mean acceleration in the medium-latency window (ANOVA p=0.096), response onset latency (ANOVA p=0.45) linear displacement (ANOVA p=0.25). (D) Quantified CoP fore-aft response, same format as (B) and (C). From left to right, mean CoP response in the short-latency window (ANOVA p=2.46e-6, regression p=1.23e-5), mean CoP response in the medium-latency window (ANOVA p=0.28), CoP response onset latency (ANOVA p=0.00103, regression p=0.00307), and CoP distance traversed (ANOVA p=0.34).

Overall, quantification of CoP responses revealed the most striking acceleration-dependent effects. Mean CoP response in the short-latency window increased significantly with acceleration (**Fig. 5D**, yellow) while the response in the medium-latency window remained unchanged (**Fig. 5D**, green). Furthermore, onset latency shifted significantly earlier with increasing acceleration (**Fig. 5D**, white, left), providing compelling evidence for acceleration-specific control of early postural responses. The distance traversed by the CoP was again small, indicating that the animals typically rode on the platform in pitch (**Fig 5D**, white, right).

### Backward pitch

We next examined responses to acceleration-controlled backward pitch perturbations (**Fig. 6A**). Two monkeys showed clear linear increases in pitch velocity during the medium-latency window, while a third monkey exhibited a similar increase in the short-latency window (**Fig. S6C**, left and middle panel). Despite these individual trends, however, overall head pitch velocity did not vary systematically with acceleration in either the short- (**Fig. 6B, yellow**) or medium-latency windows (**Fig. 6B**, green). In contrast, head fore-aft linear acceleration was systematically modulated across both windows: short-latency responses increased significantly with acceleration (**Fig. 6C**, yellow), whereas medium-latency responses decreased significantly (**Fig. 6C**, green). Additionally, the latency of the head pitch response, but not fore-aft translation, became significantly earlier with increasing acceleration (**Fig. 6B**, white, left), while acceleration onset timing remained unchanged (**Fig. 6C**, white, left). Notably, as with backward pitch of increasing velocity, backward pitch displacement fell more in the compensating window than for forward pitch and was significantly different between accelerations, but without a linear trend (**Fig. 6B**, white, right).

**Figure 6.**
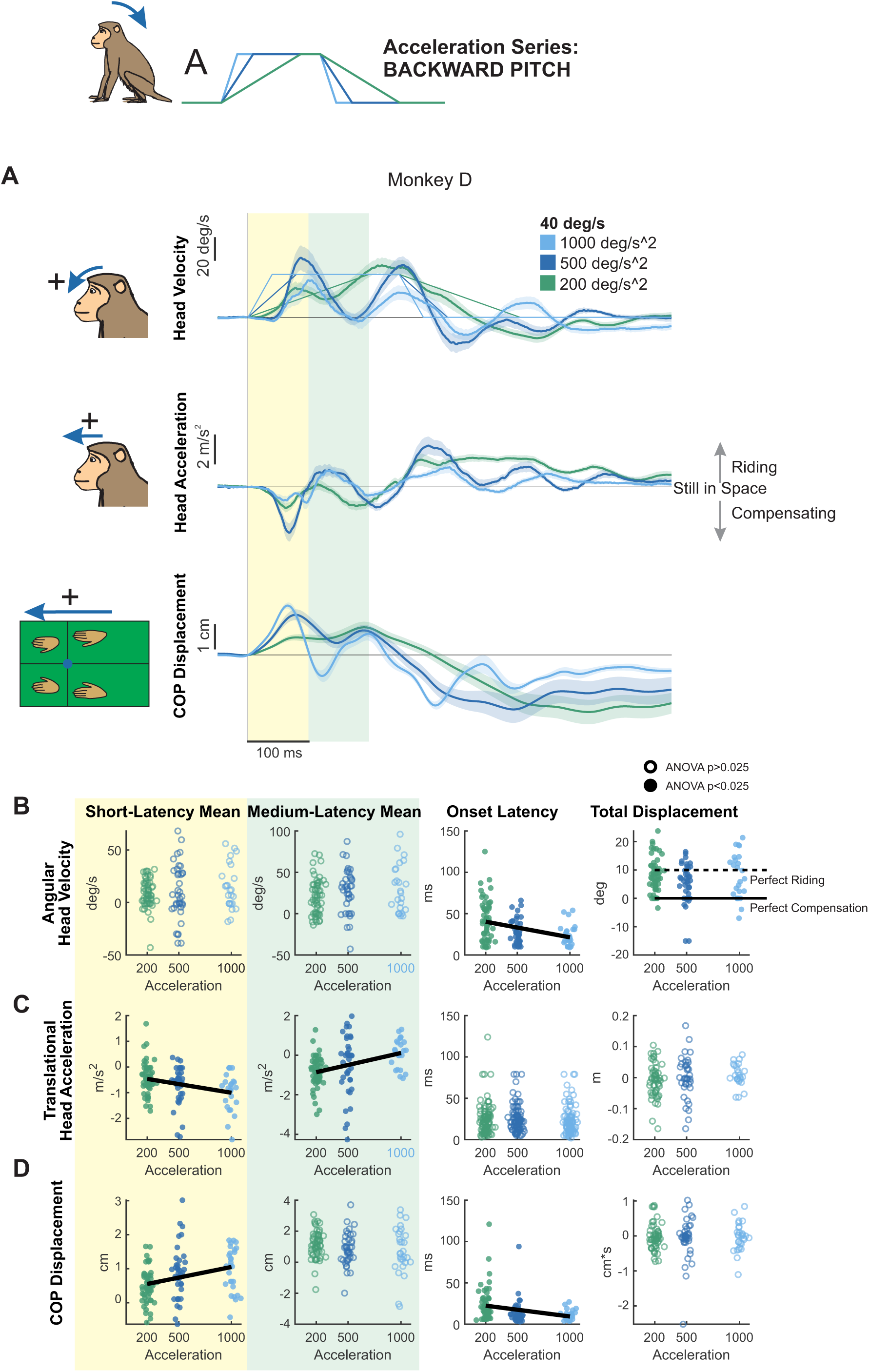
Postural responses to backward pitch perturbations of varying acceleration in the axis of the perturbation. (A) Head pitch velocity, head fore-aft acceleration, and fore-aft CoP responses of “Monkey D” to perturbations with accelerations of 200 (purple), 500 (blue), and 1000 (green) deg/s^2^ and velocity of 40 deg/s. Traces show the mean response over 20 trials and shaded regions represent the mean +/- SEM. (B) Quantified head pitch velocity response, same plotting conventions as Figure 5. From left to right: Yellow region: mean head pitch velocity in the short-latency window (ANOVA p=0.72). Green region: mean head pitch velocity in the medium-latency window (ANOVA p=0.34). White background, left: response onset latency (ANOVA p=0.000172, regression p=0.000157). White background, right: total head pitch displacement (ANOVA p=0.0184, regression p=0.63). Dotted black line indicates perfect riding response, solid black line indicates perfect compensatory response leading to no head-in-space motion. See Figure S6 for mean responses of individual monkeys. (C) Quantified head fore-aft acceleration response, same format as (B). From left to right, mean acceleration in the short-latency window (ANOVA p=0.00266, regression p=0.00115), mean acceleration in the medium-latency window (ANOVA p=0.00101, regression p=0.000209), response onset latency (ANOVA p=0.68) linear displacement (ANOVA p=0.65). (D) Quantified CoP fore-aft response, same format as (B) and (C) From left to right, mean CoP response in the short-latency window (ANOVA p=0.000540, regression p=0.000700), mean CoP response in the medium-latency window (ANOVA p=0.18), CoP response onset latency (ANOVA p=0.000753, regression p=0.00167), and CoP distance traversed (ANOVA p=0.78).

Finally, CoP responses scaled robustly with acceleration in the short-latency window (**Fig. 6D**, yellow) but remained unchanged in the medium-latency window (**Fig. 6D**, green). The onset of the CoP response also occurred significantly earlier with higher accelerations (**Fig. 6D**, white, left). Interestingly, short-latency CoP responses in the high-acceleration condition were smaller than those in other axes and CoP total distance traversed was smaller (**Fig 6D**, white, right), likely reflecting biomechanical constraints of the monkeys’ posture—forelimbs extended and hindlimbs flexed—which may limit rapid compensatory responses to backward perturbations, as well as animal-specific strategies.

### Roll axis

Using our constant-velocity protocol, we likewise recorded postural responses to roll tilts (Monkey D: **Fig. 7A**, Monkey E: **Fig. S7A**, Monkey B: **Fig. S7B**). Consistent with pitch perturbations, early head motion in roll showed acceleration-dependent modulation. Although roll velocity in the short-latency window was unaffected by acceleration (**Fig. 7B**, yellow), the onset of the roll velocity response occurred significantly earlier with increasing acceleration (**Fig. 7B**, white, left). Notably, head roll displacement was unaffected by platform acceleration and fell in the partial compensation region (**Fig 7B**, white, right). Head lateral acceleration increased significantly in the short-latency window (**Fig. 7C**, yellow), but showed no effect of acceleration in the medium-latency window (**Fig. 7C**, green). In contrast, the onset latency of lateral acceleration remained unchanged across conditions (**Fig. 7C**, white, left). Most strikingly, CoP responses exhibited robust acceleration effects: short-latency CoP responses increased markedly with higher accelerations (**Fig. 7D**, yellow), whereas medium-latency responses remained unchanged (**Fig. 7D**, green). Additionally, CoP response onset occurred significantly earlier as acceleration increased (**Fig. 7D**, white, left). Notably, this pattern contrasted with pitch perturbations, where medium-latency CoP responses decreased with increasing acceleration, highlighting axis-specific strategies—passive ‘riding’ in pitch versus active compensation in roll.

**Figure 7.**
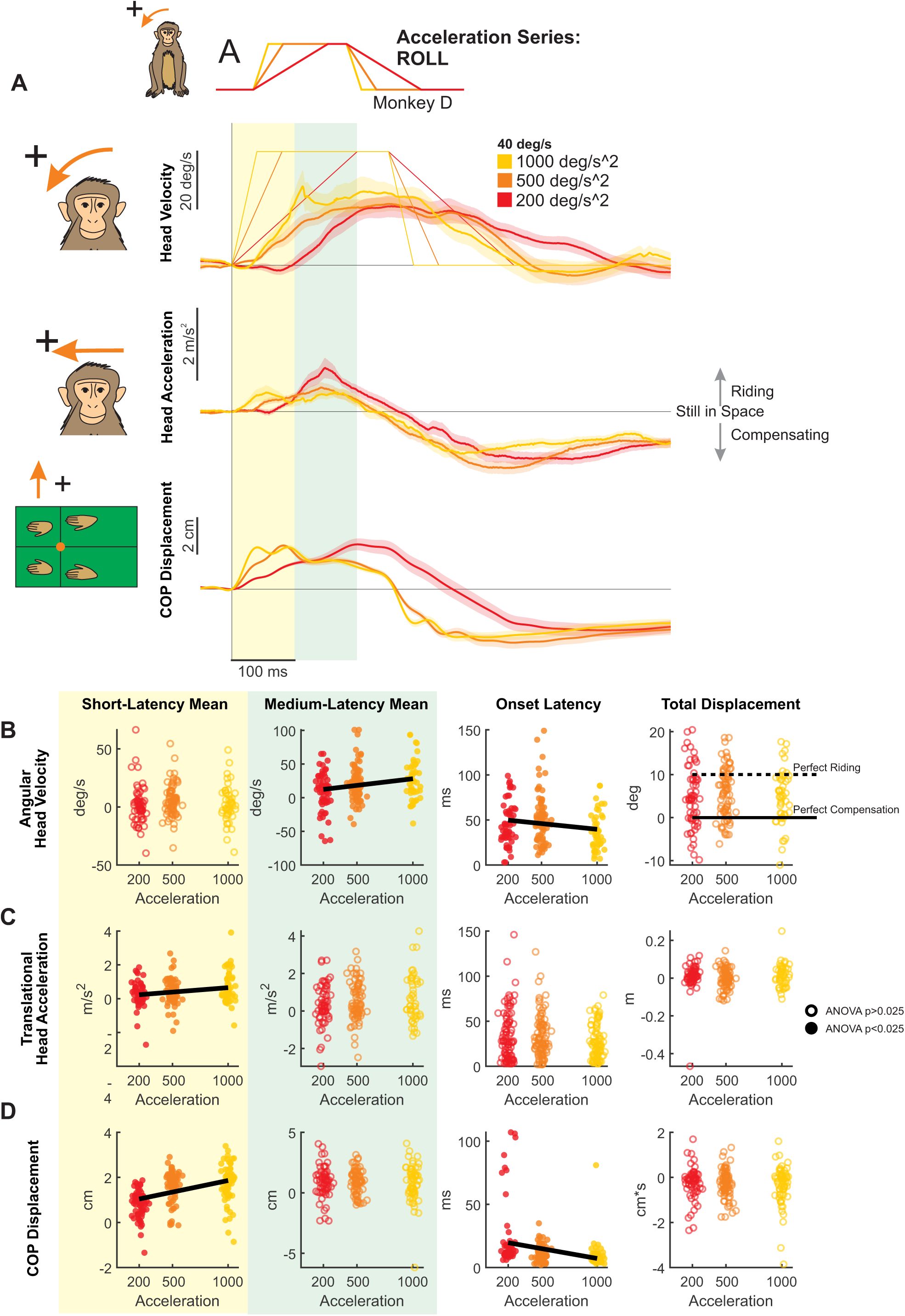
Postural responses to roll perturbations of varying acceleration in the axis of the perturbation. (A) Head roll velocity, head lateral acceleration, and lateral CoP responses of “Monkey D” to perturbations with accelerations of 200 (red), 500 (orange), and 1000 (yellow) deg/s^2^ and velocity of 40 deg/s. Traces show the mean response over 20 trials and shaded regions represent the mean +/- SEM. (B) Quantified head roll velocity response. Dots and open circles represent individual trials, pooled across monkeys, same plotting conventions as Figure 5. From left to right: Yellow region: mean head roll velocity in the short-latency window (ANOVA p=0.29). Green region: mean head roll velocity in the medium-latency window (ANOVA p=0.00383, regression p=0.00337). White background, left: response onset latency (ANOVA p=0.0112, regression p=0.0362). White background, right: total head roll displacement (ANOVA p=0.45). Dotted black line indicates perfect riding response, solid black line indicates perfect compensatory response leading to no head-in-space motion. See Figure S7 for mean responses of individual monkeys. (C) Quantified head lateral acceleration response, same format as (B). From left to right, mean acceleration in the short-latency window (ANOVA p=0.00903, regression p=0.00487), mean acceleration in the medium-latency window (ANOVA p=0.30), response onset latency (ANOVA p=0.049), and linear displacement (ANOVA p=0.27). (D) Quantified CoP lateral response, same format as (B) and (C) From left to right, mean CoP response in the short-latency window (ANOVA p=1.55e-14, regression p=6.05e-10), mean CoP response in the medium-latency window (ANOVA p=0.549), CoP response onset latency (ANOVA p=9.472e-6, regression p=0.000218) CoP distance traversed (ANOVA p=0.56).

Taken together, these findings demonstrate that angular acceleration modulates the short-latency postural response in head velocity, acceleration, and center-of-pressure dynamics—without significantly altering the magnitude of the medium-latency response—reinforcing the idea that acceleration shapes the dynamics, but not the magnitude, of early motor responses. More broadly, our results show that rhesus monkey postural responses closely mirror those of humans, supporting their use as a translational model for studying balance control. Most notably, we identify a clear dissociation between the contributions of acceleration and velocity: early (short-latency) responses are primarily driven by platform acceleration, while later (medium-latency) responses are shaped by peak velocity. In the initial phase, limb forces generated by acceleration dominate the compensatory response, whereas in the later phase, adjustments reflect steady-state motion. This dissociation provides a foundational insight into the sensory-motor basis of balance and offers a robust framework for future investigations into postural control.

## Discussion

By experimentally separating angular acceleration from velocity, we show that acceleration primarily influences the earliest (short-latency) responses, whereas velocity modulates medium-latency adjustments. The axis of rotation further shaped postural control strategies: roll tilts evoked active stabilization of the body in space, whereas pitch tilts generally elicited a compliant “ride-the-platform” response—moving along with the platform rather than stabilizing relative to space (**Fig. 8**). By separating these kinematic variables, our results resolve a key limitation of prior work and establish a primate framework that will enable future studies to directly link neural circuit activity to the temporal organization of postural control.

**Figure 8.**
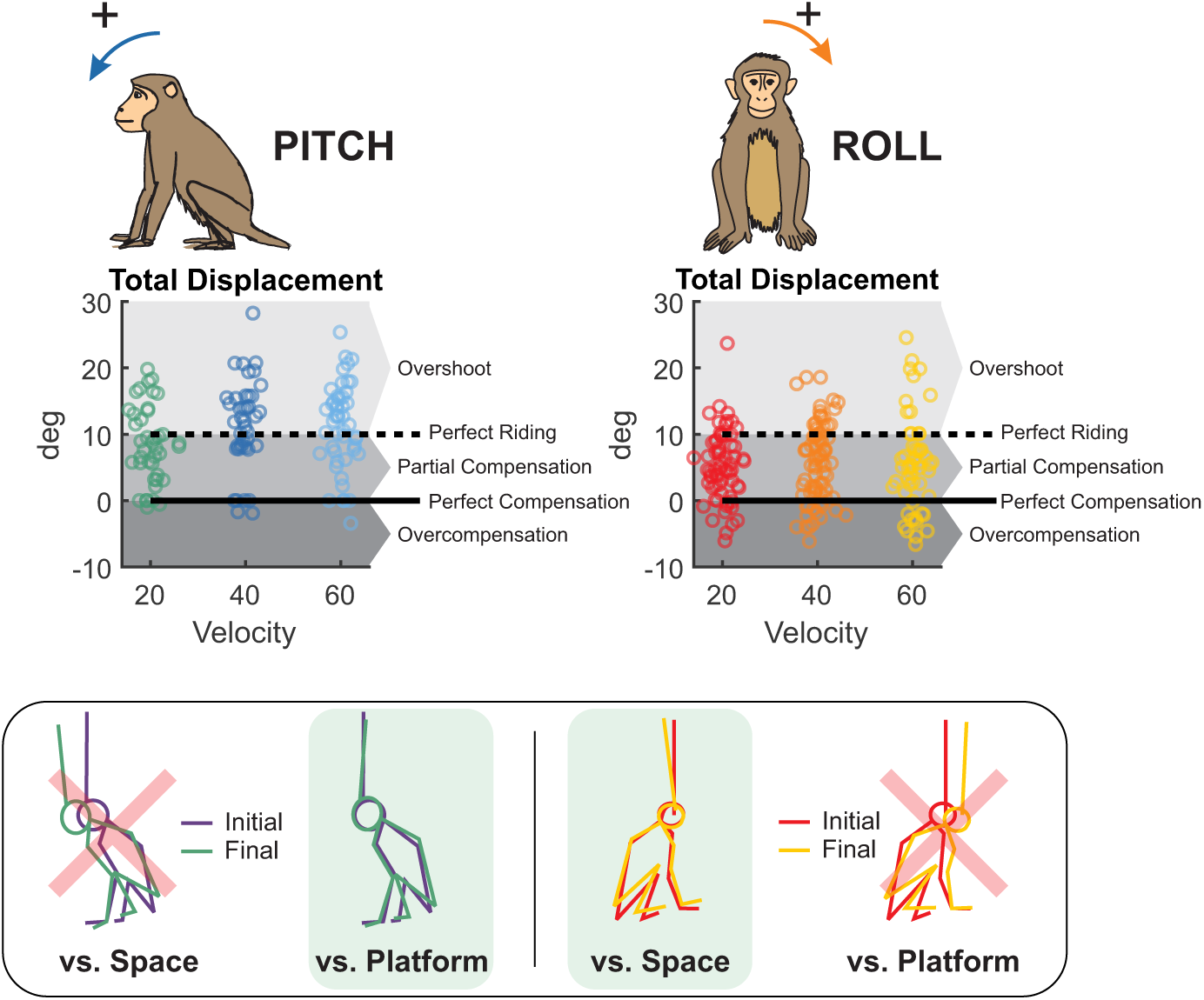
Dissociation of postural strategies across pitch and roll axes. (A) Total head angular displacement for pitch perturbations (left) and roll perturbations (right), pooled across all velocity and acceleration conditions. Individual trials are shown as dots. Dashed black line: theoretical “perfect riding” response (head displacement = platform displacement = 10°). Solid black line: theoretical “perfect compensation” (head displacement = 0°, indicating complete stabilization in space). Pitch displacements clustered at or above 10°, consistent with a passive platform-following strategy. Roll displacements fell between 0° and 10°, indicating partial active compensation. (B) Representative video frames from single trials illustrating postural strategy. Pitch (left): the body remains aligned with the tilting platform, consistent with platform-following. Roll (right): the body remains closer to vertical despite platform tilt, consistent with stabilization relative to space.

### Axis-specific biomechanical strategies

Extensive work in humans has characterized postural responses to rotational perturbations about the roll and pitch axes (Allum et al., 2008; Carpenter et al., 2001; Goodworth et al., 2023; Küng et al., 2009; Mansfield & Maki, 2009; Welch & Ting, 2009). In our nonhuman primate model, roll responses were highly symmetrical for leftward versus rightward tilts, whereas pitch responses were asymmetrical for forward versus backward tilts. This axis-dependent pattern mirrors human findings and reflects the expected symmetry along anatomically symmetric axes versus asymmetry along anatomically asymmetric axes (Allum et al., 2008; Carpenter et al., 2001; Welch & Ting, 2009).

In the roll axis, head velocity remained substantially smaller in magnitude than platform motion, indicating active stabilization of the body in space. This result is consistent with biomechanical predictions: the allowable center-of-mass excursion before balance is threatened depends on the ratio of base-of-support width to center-of-mass height in the axis of the perturbation (Forbes et al., 2018). Because this ratio is similar in humans and macaques in roll (Fryar et al., 2021; Vilensky, 1979), we predicted comparable postural strategies across species. Consistent with this prediction, rhesus monkeys exhibited roll stabilization similar to that reported in humans (Allum et al., 2008; Carpenter et al., 2001). Moreover, the only prior study examining roll perturbations in monkeys—using continuous pseudorandom stimulation—also found postural sway to be smaller than platform displacement (Thompson et al., 2016), supporting the robustness of roll stabilization across perturbation paradigms.

In contrast, pitch tilts elicited a more compliant postural strategy in which animals largely moved with the platform, tolerating larger center-of-mass displacements before generating corrective torques. This behavior likely reflects the elongated anteroposterior base of support produced by coordinated fore- and hindlimb stabilization, which mechanically increases tolerance to pitch perturbations. Strikingly, even the most dynamic pitch perturbations tested here (1000 deg/s²) failed to elicit strong stabilizing responses, paralleling findings in quadrupedal cats (Jacobs & Macpherson, 1996; Macpherson et al., 2007). Humans, in contrast, exhibit strong pitch stabilization at much lower accelerations (260 deg/s²) (Allum et al., 2008; Carpenter et al., 2001; Horak et al., 2016), but adopt cat- and monkey-like pitch strategies when assuming a quadrupedal stance (Dunbar et al., 1986; Macpherson et al., 2007). Together, these comparisons highlight how body morphology and base-of-support geometry shape axis-specific postural strategies while preserving common vestibular control principles across species.

### Balance responses to rotational platform perturbations: dissociating acceleration from velocity

Prior studies using rotational platform perturbations have consistently confounded angular acceleration and peak velocity, precluding clear attribution of their relative contributions (reviewed in Forbes et al., 2020). By contrast, prior studies focused on postural responses to translational platform perturbations have successfully dissociated the effects of acceleration versus velocity. Specifically, acceleration has been shown to dominate short-latency responses to translational perturbations, while velocity primarily influences medium-latency components (Allum et al., 1994; Allum & Pfaltz, 1985; Bloem et al., 2002; Diener et al., 1988; Horak et al., 1990; Inglis & Macpherson, 1995; Maki & Ostrovski, 1993; Welch & Ting, 2009). Our study extends this dissociation to rotational perturbations, demonstrating that short-latency responses are primarily acceleration-driven, whereas medium-latency responses scale with velocity. Notably, this parallel across perturbation types emerges despite fundamental differences in destabilizing mechanics—translation destabilizes posture through trunk inertia, whereas rotational tilt can initially be mechanically stabilizing (Lippi et al., 2023; reviewed in Mergner et al., 2009). Because each monkey contributed many repeated trials across conditions, the within-subject design enabled robust quantification of response dynamics, and the dissociation between acceleration-dependent short-latency responses and velocity-dependent medium-latency responses was consistent across animals.

Overall, the present study establishes that, paralleling translational perturbations, stability during dynamic tilts follows a systematic and dissociable temporal sequence. Such a sequence requires continuous estimation and correction of body orientation relative to both the support surface and gravity, implemented through precisely timed and scaled joint torques (Missen et al., 2023; Peterka, 2018). By orthogonalizing angular acceleration and peak velocity, our design provides the first clear demonstration of this dissociation in rotational postural control, offering a methodological framework that can now be extended to probe the neural circuits that implement these computations.

### Vestibular contributions revealed by head motion

Most postural studies focus on CoP, limb kinematics, or EMG, yet head motion provides a direct measure of the stimulus driving vestibular afferents and the resulting motor response. Because vestibular afferents encode head motion itself (reviewed in Cullen 2019), voluntary head movements simultaneously represent motor outputs and sensory signals for self-motion encoding. Consequently, vestibular signals encoding head-centered motion must be transformed into body-centered coordinates to generate appropriate postural adjustments (Keshner & Dhaher, 2008; Mergner, 2007; Peterka, 2002). Because platform tilts move the entire support surface, postural responses reflect integrated vestibular and somatosensory feedback: head kinematics characterize the vestibular stimulus, whereas postural motion reflects multisensory feedback integration. For this reason, we quantified head kinematics together with CoP dynamics to capture both the vestibular stimulus and the resulting whole-body postural response.

In the present study, direct measurements of six-dimensional head motion revealed that, like CoP, evoked head motion was influenced by both acceleration and velocity. Notably, compared to the few prior human studies that have measured head motion responses, head motion in monkeys began substantially earlier—within ∼60 ms for transient roll stimuli and as early as ∼20 ms for pitch—versus ∼80–120 ms in humans (Allum, 1992; Carpenter et al., 2001; Horak & Macpherson, 1996). This difference in onset timing likely reflects a combination of biomechanical and stimulus factors, including shorter limb lengths and our use of higher angular accelerations (200–1000 deg/s² versus 8–260 deg/s² in human studies). Consistent with this interpretation, at the lowest acceleration tested, roll latencies approached values previously reported in humans, reinforcing the role of acceleration in determining response onset.

It is noteworthy that previous work in patients with proprioceptive deficits (Carpenter et al., 2001) and vestibular loss (Horak et al., 2016), as well as studies using galvanic vestibular stimulation (Horak & Hlavacka, 2002), suggests that short-latency responses are dominated by proprioceptive inputs, with vestibular contributions emerging at longer latencies. For translational perturbations, otolith afferents encode linear acceleration (or jerk under highly dynamic conditions; Jamali et al., 2013, 2019), limiting velocity-dependent effects. In contrast, during rotational perturbations, semicircular canals encode angular velocity—and acceleration under dynamic conditions (Jamali et al., 2016; Sadeghi et al., 2007), allowing both kinematic variables to shape postural responses. Nevertheless, across perturbation types, platform motion ultimately induces whole-body sway, requiring continuous monitoring of body motion relative to gravity and the support surface (Mergner et al., 2009; Peterka, 2018). The conserved sequence of acceleration- and velocity-modulated response phases observed here, corresponding to previously reported EMG bursts (Diener et al., 1983; Dunbar et al., 1986), supports a general control mechanism regulating postural dynamics across axes.

### Rhesus macaques as a platform for circuit-level dissection of balance control

Rhesus macaques provide a valuable translational bridge between classical animal models and human studies of postural control. While cats have traditionally served as models for postural reflexes, macaques share over 97% genomic similarity with humans (Gibbs et al., 2007), exhibit similar head-movement statistics during natural behaviors (Carriot et al., 2017), and—unlike digitigrade cats—are plantigrade, bearing weight across the entire foot. This posture imposes biomechanical demands for standing balance that closely resemble those in humans. Consistent with this alignment, rhesus monkey CoP responses—particularly in roll—closely parallel those reported in humans (Dunbar et al., 1986), despite earlier response onsets consistent with body scaling, indicating conserved neural control strategies across primates. In contrast, differences in body plan shape responses to pitch perturbations, whereas roll responses remain highly similar across species. Together, these findings establish behavioral benchmarks for interpreting postural control in the macaque model. Because this model permits invasive recordings and causal circuit manipulations not feasible in humans, future studies can determine how neural population activity contributes to distinct temporal phases of the postural response and evaluate potential vestibular prosthetic interventions (Boutros et al., 2019; Wiboonsaksakul et al., 2022, 2023).

Our experiments focused on transient perturbations, leaving open whether acceleration- and velocity-dependent control generalizes to prolonged or stochastic disturbances encountered during natural behavior. Moreover, although our approach isolates the kinematic variables shaping postural responses, the sensory–motor circuits implementing these axis-dependent strategies remain unknown. Future studies combining this perturbation framework with neural recordings or causal manipulations in rhesus monkeys will be essential for linking circuit dynamics to the temporal phases of balance control and guiding vestibular prosthetic strategies.

## Supporting information

All Supplemental Material

## Acknowledgements

We would like to thank the entire Cullen Lab especially Dale Roberts, Walter Kucharski, and Balazs Vagvolgyi for their technical support; and Pum Wiboonsaksakul, Robyn Mildren, Lex Gómez, Skyler Thomas, Chenhao Bao, and Oliver Stanley for their advice and comments on the manuscript and figures. We would also like to thank the animal care and veterinary staff as well as our animal subjects. This research was supported by grants R01DC002390 & R01DC018061 from the National Institutes of Health (K.E.C.), as well as Ruth L. Kirchstein National Research Service Award F31DC020390 (O.L.B.).

## Author Contributions

**O.M.E.L.B.:** Conceptualization, Methodology, Software, Formal Analysis, Investigation, Data Curation, Visualization, Writing – Original Draft, Writing – Review & Editing. **B.A.R.:** Investigation, Data Curation, Writing – Review & Editing. **K.E.C.:** Conceptualization, Resources, Writing – Review & Editing, Supervision, Funding Acquisition.

## Data Availability

Data and analysis code are available from the corresponding author upon reasonable request.

