## Supplementary material for "Acceleration and Velocity Dissociate Temporal Phases of Postural Control in Rhesus Macaques": All Supplemental Material

**Supplementary Figure Legends**

**Figure S1**. Postural responses to forward pitch perturbations of varying peak velocity in the axis of the perturbation. (A) Head pitch velocity, head fore-aft acceleration, and fore-aft CoP responses of “Monkey E” to perturbations with peak velocities of 20 (purple), 40 (blue), and 60 (green) deg/s and acceleration of 500 deg/s$^2$. Traces show the mean response over 20 trials and shaded regions represent the mean +/- SEM. (B) same as (A) for “Monkey B”. (C) Quantified head pitch velocity response by animal. Markers represent mean response of individual animals, with error bars representing SEM. Filled markers indicate statistical significance (ANOVA $p<0.025$) of comparison across monkeys independent of trial condition. Open markers indicate lack of statistical significance (ANOVA $p>0.025$). Gray bars show mean across animals. From left to right: Yellow region: mean head pitch velocity in the short-latency window (ANOVA p=9.67e-13). Green region: mean head pitch velocity in the medium-latency window (ANOVA p=6.144e-5). White background, left: response onset latency (ANOVA p=0.027). White background, right: total head pitch displacement (ANOVA p=0.46). (D) Quantified head fore-aft acceleration response, same format as (B). From left to right, mean acceleration in the short-latency window (ANOVA p=0.000189), mean acceleration in the medium-latency window (ANOVA p=0.00975), response onset latency (ANOVA p=0.29), and linear displacement (ANOVA p=1.74e-8). (E) Quantified CoP fore-aft response, same format as (B) and (C). From left to right, mean CoP response in the short-latency window (ANOVA p=0.21), mean CoP response in the medium-latency window (ANOVA p=2.60e-7), CoP response onset latency (ANOVA p=4.34e-8), and CoP distance traversed (ANOVA p=0.31).

**Figure S2**. Postural responses to backward pitch perturbations of varying peak velocity in the axis of the perturbation. (A) Head pitch velocity, head fore-aft acceleration, and fore-aft CoP responses of “Monkey E” to perturbations with peak velocities of 20 (purple), 40 (blue), and 60 (green) deg/s and acceleration of 500 deg/s$^2$. Traces show the mean response over 20 trials and shaded regions represent the mean +/- SEM. (B) same as (A) for “Monkey B”. (C) Quantified head pitch velocity response by animal, same plotting conventions as Figure S1. From left to right: Yellow region: mean head pitch velocity in the short-latency window (ANOVA p=5.55e-13). Green region: mean head pitch velocity in the medium-latency window (ANOVA p=0.0251). White background, left: response onset latency (ANOVA p=0.21). White background, right: total head pitch displacement (ANOVA p=4.13e-6). (D) Quantified head fore-aft acceleration response, same format as (B). From left to right, mean acceleration in the short-latency window (ANOVA p=0.00354), mean acceleration in the medium-latency window (ANOVA p=4.84e-12), response onset latency (ANOVA p=0.71), and linear displacement (ANOVA p=3.10e-11). (E) Quantified CoP fore-aft response, same format as (B) and (C). From left to right, mean CoP response in the short-latency window (ANOVA p=0.41), mean CoP response in the medium-latency window (ANOVA p=0.0185), CoP response onset latency (ANOVA p=0.64), and CoP distance traversed (ANOVA p=9.03e-6).

**Figure S3**. Symmetry of pitch vs. roll perturbation responses. (A) Head pitch velocity, fore-aft acceleration, and fore-aft CoP response to forward (solid) vs backward (dotted) perturbations, averaged across all monkeys. (B) Head roll velocity, lateral acceleration, and lateral CoP response to roll perturbations. (C) Symmetry index for pitch (blue) vs roll (orange) perturbations in velocity, acceleration, and CoP response. Roll (head velocity p=0.12, head acceleration p=0.14, CoP response p=0.49) had consistently symmetrical responses, whereas pitch (head velocity p=0.18, head acceleration p=0.0216, CoP response p=2.82e-7) had a mix of symmetrical and asymmetrical.

**Figure S4**. Postural responses to roll perturbations of varying peak velocity in the axis of the perturbation. (A) Head roll velocity, head lateral acceleration, and lateral CoP responses of “Monkey E” to perturbations with peak velocities of 20 (red), 40 (orange), and 60 (yellow) deg/s and acceleration of 500 deg/s$^2$. Traces show the mean response over 20 trials and shaded regions represent the mean +/- SEM. (B) same as (A) for “Monkey B”. (C) Quantified head roll velocity response by animal, same plotting conventions as Figure S1. From left to right: Yellow region: mean head roll velocity in the short-latency window (ANOVA p=0.35). Green region: mean head roll velocity in the medium-latency window (ANOVA p=8.07e-9). White background, left: response onset latency (ANOVA p=0.002). White background, right: total head roll displacement (ANOVA p=1.380e-5). (D) Quantified head lateral acceleration response, same format as (B). From left to right, mean acceleration in the short-latency window (ANOVA p=0.000950), mean acceleration in the medium-latency window (ANOVA p=0.0166), response onset latency (ANOVA p=0.13), and linear displacement (ANOVA p=0.19). (E) Quantified CoP lateral response, same format as (B) and (C). From left to right, mean CoP response in the short-latency window (ANOVA p=0.000145), mean CoP response in the medium-latency window (ANOVA p=0.000568), CoP response onset latency (ANOVA p=0.00146), and CoP distance traversed (ANOVA p=0.42).

**Figure S5**. Postural responses to forward pitch perturbations of varying acceleration in the axis of the perturbation. (A) Head pitch velocity, head fore-aft acceleration, and fore-aft CoP responses of “Monkey E” to perturbations with accelerations of 200 (purple), 500 (blue), and 1000 (green) deg/s$^2$ and velocity of 40 deg/s. Traces show the mean response over 20 trials and shaded regions represent the mean +/- SEM. (B) same as (A) for “Monkey B”. (C) Quantified head pitch velocity response by animal, same plotting conventions as Figure S1. From left to right: Yellow region: mean head pitch velocity in the short-latency window (ANOVA p=4.53e-8). Green region: mean head pitch velocity in the medium-latency window (ANOVA p=0.44). White background, left: response onset latency (ANOVA p=0.054). White background, right: total head pitch displacement (ANOVA p=0.69). (D) Quantified head fore-aft acceleration response, same format as (B). From left to right, mean acceleration in the short-latency window (ANOVA p=0.00137), mean acceleration in the medium-latency window (ANOVA p=5.84e-6), response onset latency (ANOVA p=0.17), and linear displacement (ANOVA p=0.0602). (E) Quantified CoP fore-aft response, same format as (B) and (C). From left to right, mean CoP response in the short-latency window (ANOVA p=0.14), mean CoP response in the medium-latency window (ANOVA p=0.000260), CoP response onset latency (ANOVA p=4.58e-5), and CoP distance traversed (ANOVA p=0.23).

**Figure S6**. Postural responses to backward pitch perturbations of varying acceleration in the axis of the perturbation. (A) Head pitch velocity, head fore-aft acceleration, and fore-aft CoP responses of “Monkey E” to perturbations with accelerations of 200 (purple), 500 (blue), and 1000 (green) deg/s$^2$ and velocity of 40 deg/s. Traces show the mean response over 20 trials and shaded regions represent the mean +/- SEM. (B) same as (A) for "Monkey B". (C) Quantified head pitch velocity response by animal, same plotting conventions as Figure S1. From left to right: Yellow region: mean head pitch velocity in the short-latency window (ANOVA p=1.61e-10). Green region: mean head pitch velocity in the medium-latency window (ANOVA p=6.19e-5). White background, left: response onset latency (ANOVA p=0.12). White background, right: total head pitch displacement (ANOVA p=1.01e-9). (D) Quantified head fore-aft acceleration response, same format as (B). From left to right, mean acceleration in the short-latency window (ANOVA p=0.13), mean acceleration in the medium-latency window (ANOVA p=4.47e-5), response onset latency (ANOVA p=0.84), and linear displacement (ANOVA p=1.36e-7). (E) Quantified CoP fore-aft response, same format as (B) and (C). From left to right, mean CoP response in the short-latency window (ANOVA p=0.097), mean CoP response in the medium-latency window (ANOVA p=0.0436), CoP response onset latency (ANOVA p=0.00441), and CoP distance traversed (ANOVA p=4.97e-9).

**Figure S7**. Postural responses to roll perturbations of varying acceleration in the axis of the perturbation. (A) Head roll velocity, head lateral acceleration, and lateral CoP responses of "Monkey E" to perturbations with accelerations of 200 (red), 500 (orange), and 1000 (yellow) deg/$s^2$ and velocity of 40 deg/s. Traces show the mean response over 20 trials and shaded regions represent the mean +/- SEM. (B) same as (A) for "Monkey B". (C) Quantified head roll velocity response by animal, same plotting conventions as Figure S1. From left to right: Yellow region: mean head roll velocity in the short-latency window (ANOVA p=0.38). Green region: mean head roll velocity in the medium-latency window (ANOVA p=0.0114). White background, left: response onset latency (ANOVA p=0.00531). White background, right: total head roll displacement (ANOVA p=2.27e-5). (D) Quantified head lateral acceleration response, same format as (B). From left to right, mean acceleration in the short-latency window (ANOVA p=0.000162), mean acceleration in the medium-latency window (ANOVA p=0.94), response onset latency (ANOVA p=0.43), and linear displacement (ANOVA p=0.027). (E) Quantified CoP lateral response, same format as (B) and (C). From left to right, mean CoP response in the short-latency window (ANOVA p=0.000162), mean CoP response in the medium-latency window (ANOVA p=2.53e-7), CoP response onset latency (ANOVA p=0.035), and CoP distance traversed (ANOVA p=0.081).

**Table S1**. Variables and values used in the dual-pendulum kinematic model of passive motion.

### Figure S1

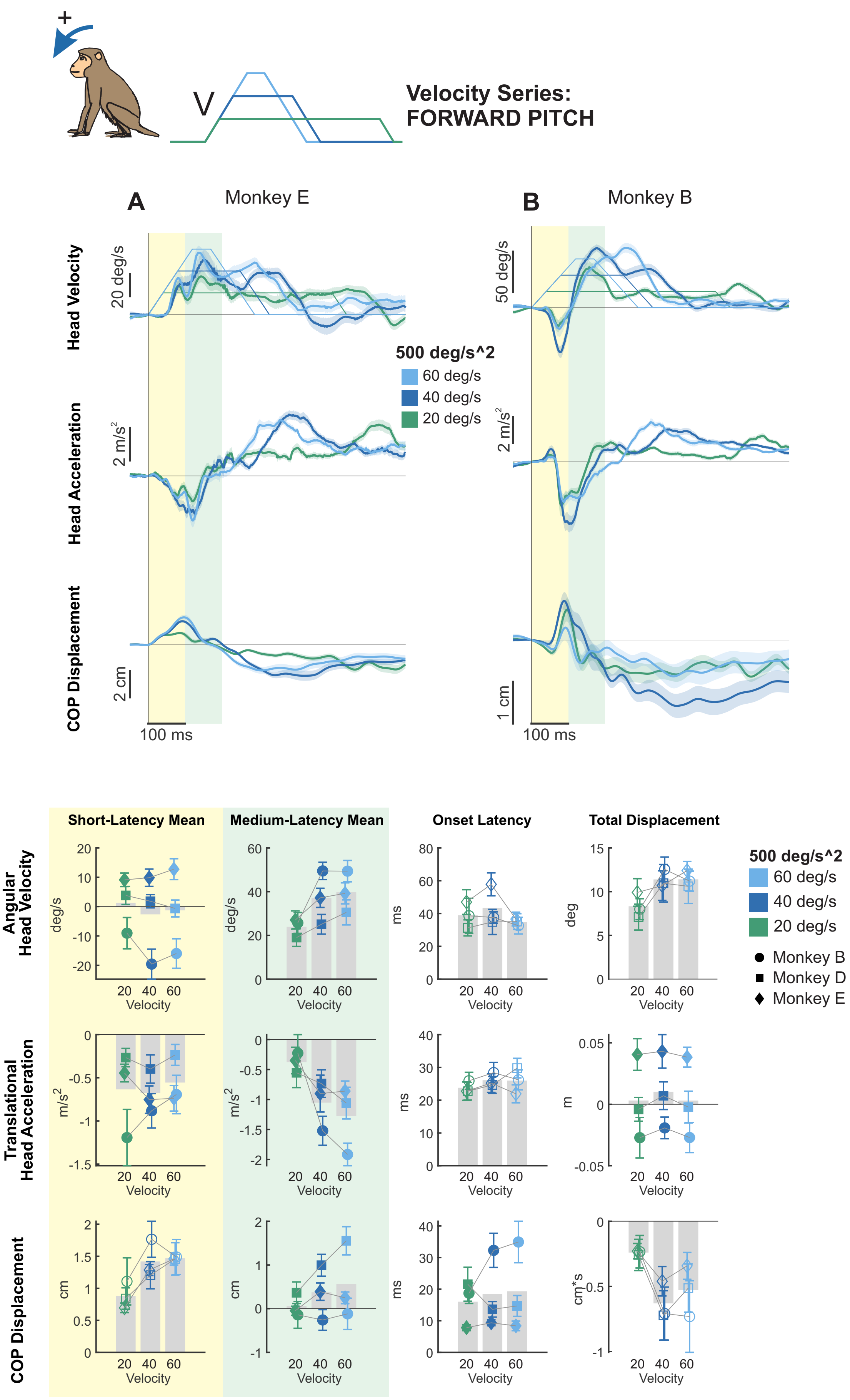
+
V
Velocity Series:
FORWARD PITCH
A
Monkey E
B
Monkey B
Head Velocity
Head Acceleration
COP Displacement
20 deg/s
50 deg/s
2 m/s^2
2 cm
1 cm
100 ms
500 deg/s^2
60 deg/s
40 deg/s
20 deg/s
C
D
E
Short-Latency Mean
Medium-Latency Mean
Onset Latency
Total Displacement
Angular Head Velocity
Translational Head Acceleration
COP Displacement
deg/s
m/s^2
cm
ms
deg
m
cm*s
Velocity
20 40 60
Monkey B
Monkey D
Monkey E

### Figure S2

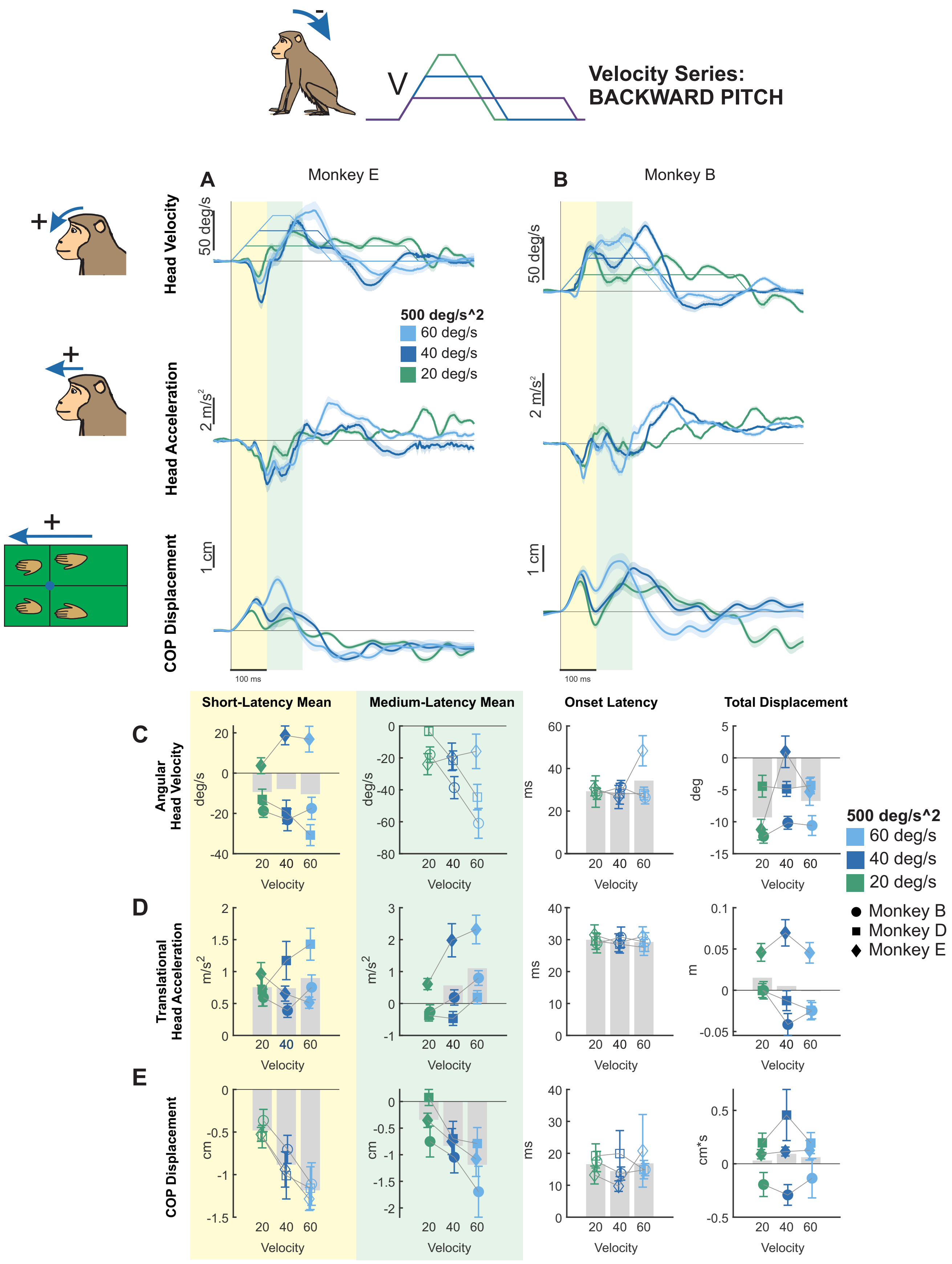

Velocity Series:
BACKWARD PITCH
V
A
Monkey E
B
Monkey B
Head Velocity
Head Acceleration
COP Displacement
50 deg/s
2 m/s²
1 cm
100 ms
500 deg/s^2
60 deg/s
40 deg/s
20 deg/s
Short-Latency Mean
Medium-Latency Mean
Onset Latency
Total Displacement
C
Angular Head Velocity
D
Translational Head Acceleration
E
COP Displacement
Velocity
deg/s
m/s²
cm
ms
deg
m
cm*s
Monkey B
Monkey D
Monkey E

### Figure S3

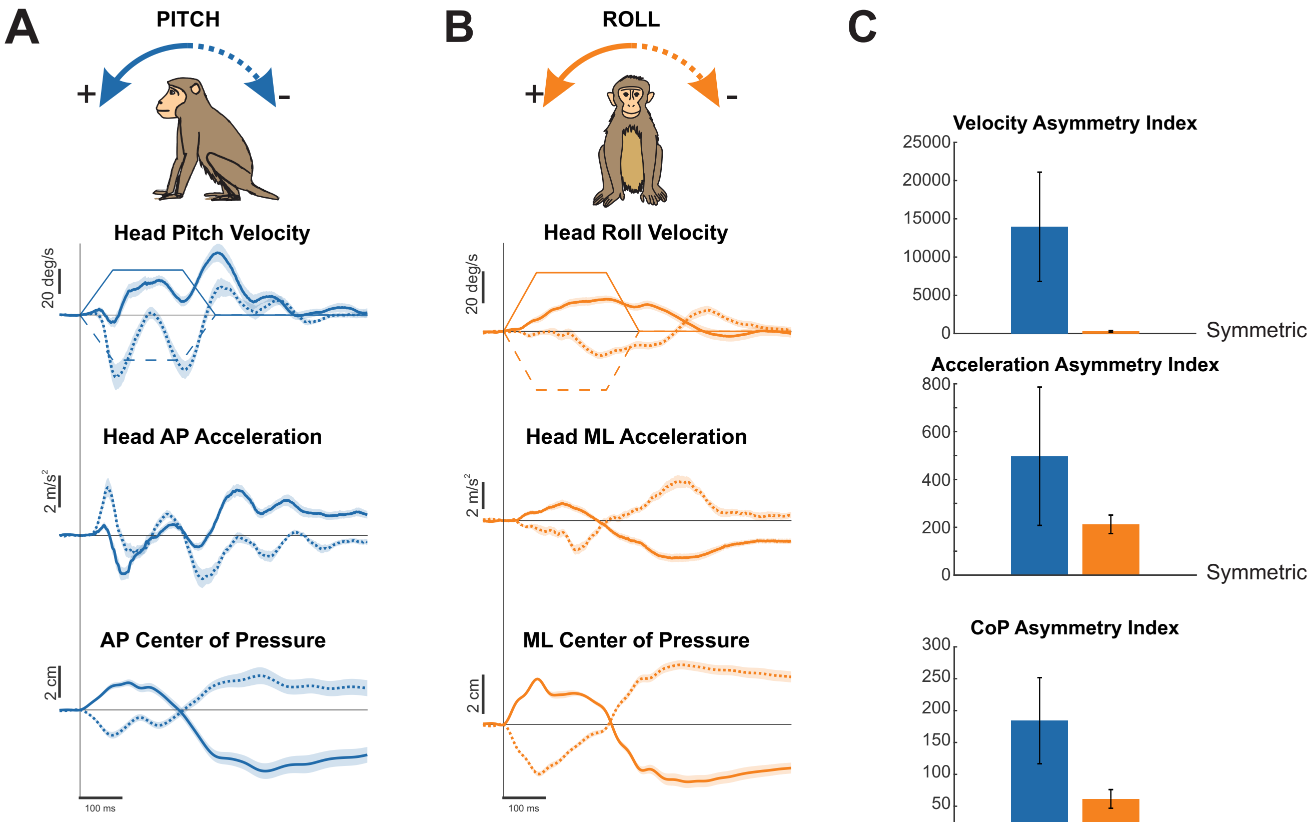
A
PITCH
+
-
Head Pitch Velocity
20 deg/s
Head AP Acceleration
2 m/s²
AP Center of Pressure
2 cm
100 ms
B
ROLL
+
-
Head Roll Velocity
20 deg/s
Head ML Acceleration
2 m/s²
ML Center of Pressure
2 cm
100 ms
C
Velocity Asymmetry Index
25000
20000
15000
10000
5000
0
Symmetric
Acceleration Asymmetry Index
800
600
400
200
0
Symmetric
CoP Asymmetry Index
300
250
200
150
100
50
0
Symmetric

### Figure S4

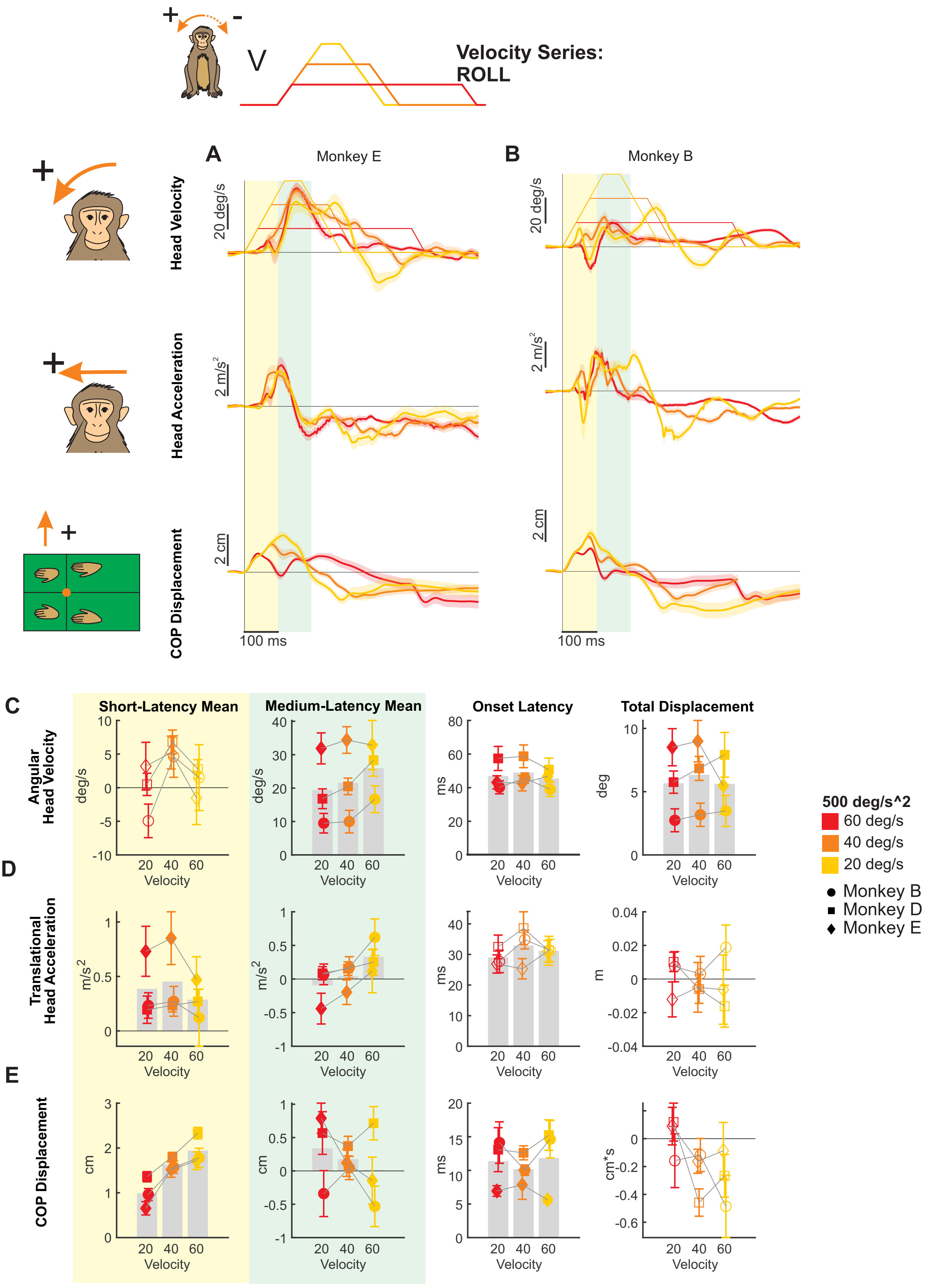

Velocity Series: ROLL
V
A
Monkey E
B
Monkey B
Head Velocity
20 deg/s
Head Acceleration
2 m/s2
COP Displacement
2 cm
100 ms
C
D
E
Short-Latency Mean
Medium-Latency Mean
Onset Latency
Total Displacement
Angular Head Velocity
Translational Head Acceleration
COP Displacement
deg/s
m/s2
cm
ms
deg
m
cm*s
Velocity
500 deg/s^2
60 deg/s
40 deg/s
20 deg/s
Monkey B
Monkey D
Monkey E

### Figure S5

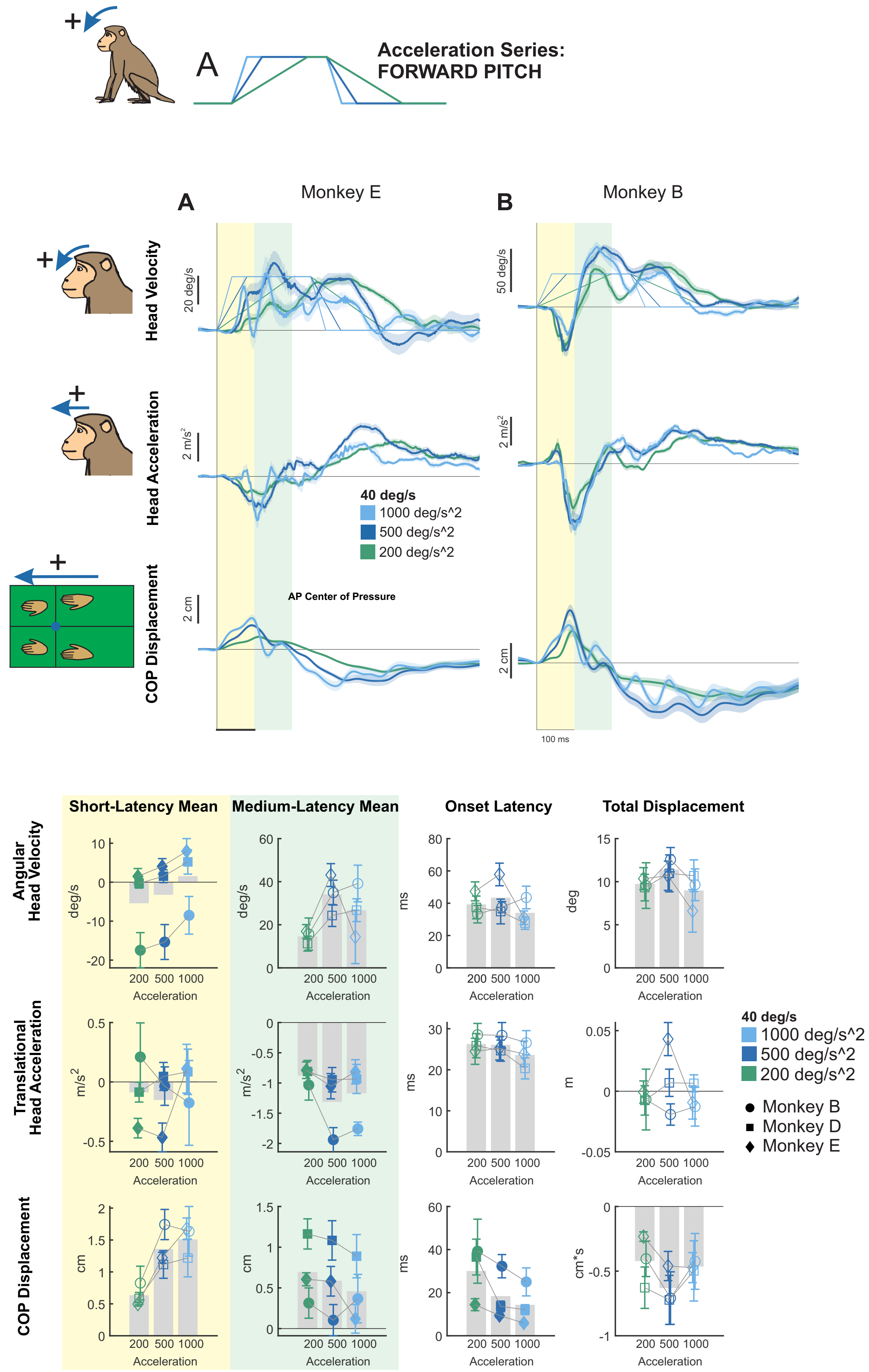

A
Acceleration Series:
FORWARD PITCH
A
Monkey E
B
Monkey B
Head Velocity
Head Acceleration
COP Displacement
20 deg/s
50 deg/s
2 m/s^2
2 cm
40 deg/s
1000 deg/s^2
500 deg/s^2
200 deg/s^2
AP Center of Pressure
100 ms
C
D
E
Short-Latency Mean
Medium-Latency Mean
Onset Latency
Total Displacement
Angular Head Velocity
Translational Head Acceleration
COP Displacement
deg/s
m/s^2
cm
ms
deg
m
cm*s
Acceleration
200 500 1000
Monkey B
Monkey D
Monkey E

### Figure S6

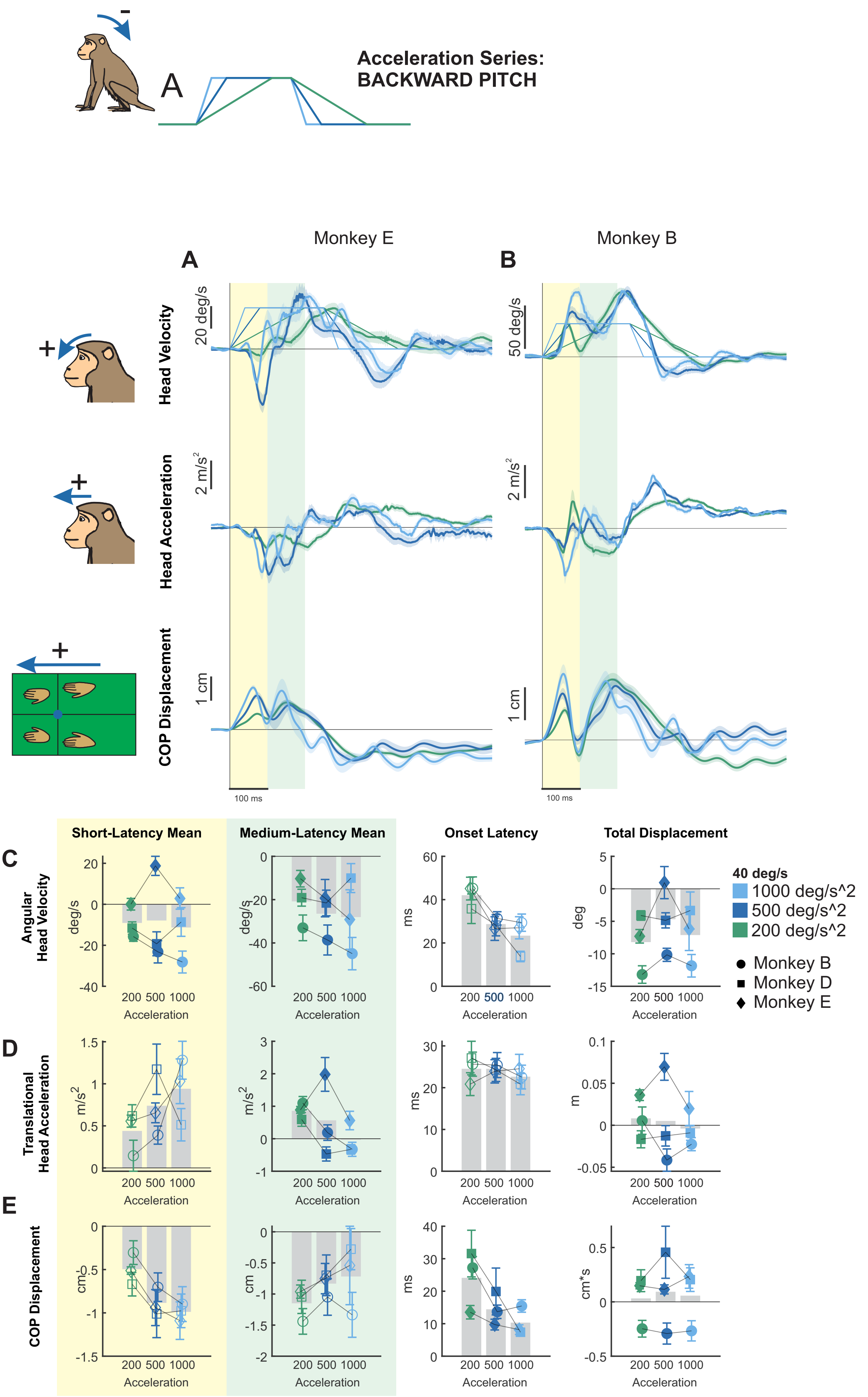

Acceleration Series:
BACKWARD PITCH
A
Monkey E
Monkey B
A
B
Head Velocity
Head Acceleration
COP Displacement
20 deg/s
50 deg/s
2 m/s^2
1 cm
100 ms
Short-Latency Mean
Medium-Latency Mean
Onset Latency
Total Displacement
C
Angular Head Velocity
D
Translational Head Acceleration
E
COP Displacement
Acceleration
200 500 1000
deg/s
ms
deg
m/s^2
m
cm
cm*s
40 deg/s
1000 deg/s^2
500 deg/s^2
200 deg/s^2
Monkey B
Monkey D
Monkey E

### Figure S7

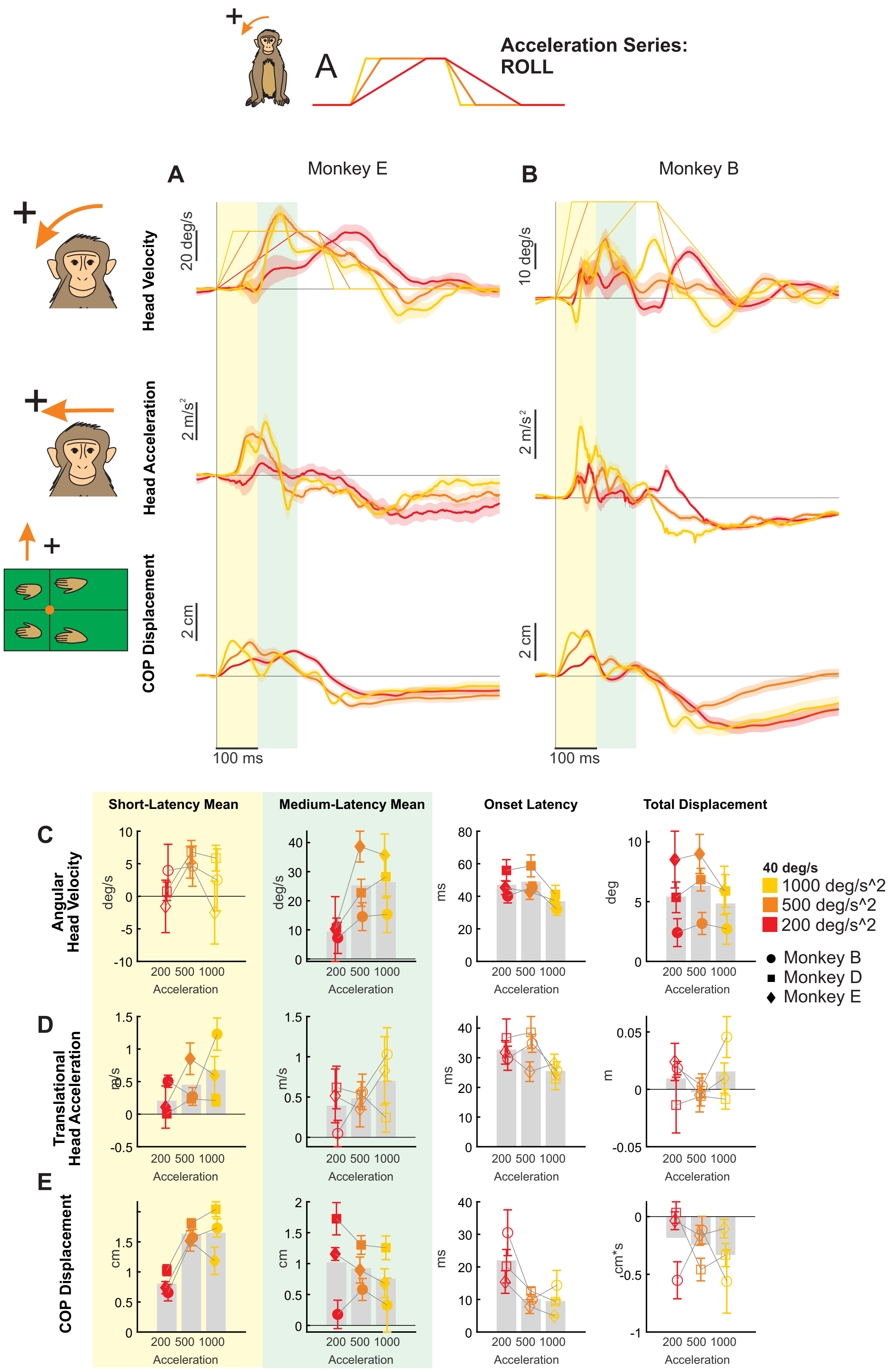

A
Acceleration Series:
ROLL
A
Monkey E
B
Monkey B
Head Velocity
Head Acceleration
COP Displacement
20 deg/s
10 deg/s
2 m/s²
2 cm
100 ms
Short-Latency Mean
Medium-Latency Mean
Onset Latency
Total Displacement
C
Angular Head Velocity
D
Translational Head Acceleration
E
COP Displacement
deg/s
m/s
cm
ms
deg
m
cm*s
Acceleration
200 500 1000
40 deg/s
1000 deg/s^2
500 deg/s^2
200 deg/s^2
Monkey B
Monkey D
Monkey E

**Supplementary Table S1**

| Variable name | Parameter | Value |
|---|---|---|
| $m_1$ | Mass of trunk and limb segment | 8.0 kg |
| $m_2$ | Mass of head | 0.5 kg |
| $l_1$ | Trunk center of mass distance | 0.35 m |
| $l_2$ | Head center of mass distance | 0.1 m |
| $I_1$ | Trunk inertia | 0.25 kg*m^2 |
| $I_2$ | Head inertia | 0.0025 kg*m^2 |
| *k_platform* | Platform-trunk coupling stiffness (“stiff”) | 10 N*m/rad |
| *k_platform* | Platform-trunk coupling stiffness (“floppy”) | 1 N*m/rad |
| *k_neck* | Trunk-head coupling stiffness (“stiff”) | 100 N*m/rad |
| *k_neck* | Trunk-head coupling stiffness (“floppy”) | 10 N*m/rad |
| *c_platform* | Platform-trunk damping | 2 N*m*s/rad |
| *c_neck* | Trunk-head damping | 0.1 N*m*s/rad |
